# Balancing growth amidst salinity stress – lifestyle perspectives from the extremophyte model *Schrenkiella parvula*

**DOI:** 10.1101/2021.08.27.457575

**Authors:** Kieu-Nga Tran, Pramod Pantha, Guannan Wang, Narender Kumar, Chathura Wijesinghege, Dong-Ha Oh, Nick Duppen, Hongfei Li, Hyewon Hong, John C. Johnson, Ross Kelt, Megan G. Matherne, Ashley Clement, David Tran, Colt Crain, Prava Adhikari, Yanxia Zhang, Maryam Foroozani, Guido Sessa, John C. Larkin, Aaron P. Smith, David Longstreth, Patrick Finnegan, Christa Testerink, Simon Barak, Maheshi Dassanayake

## Abstract

*Schrenkiella parvula*, a leading extremophyte model in Brassicaceae, can grow and complete its life cycle under multiple environmental stresses, including high salinity. While foundational genomic resources have been created for *S. parvula*, a comprehensive physiological or structural characterization of its salt stress responses is absent. We aimed to identify the influential traits that lead to stress-resilient growth of this species. We examined salt-induced changes in the physiology and anatomy of *S. parvula* throughout its lifecycle across multiple tissues. We found that *S. parvula* maintains or even exhibits enhanced growth during various developmental stages at salt stress levels known to inhibit growth in Arabidopsis and most crops. The resilient growth of *S. parvula* was associated with key traits that synergistically allow continued primary root growth, expansion of xylem vessels across the root-shoot continuum, and a high capacity to maintain tissue water levels by developing larger and thicker leaves while facilitating continued photosynthesis during salt stress. These traits at the vegetative phase were followed by a successful transition to the reproductive phase via early flowering, development of larger siliques, and production of viable seeds during salt stress. Additionally, the success of self-fertilization during early flowering stages was dependent on salt-induced filament elongation in flowers that aborted in the absence of salt. Our results suggest that the maintenance of leaf water status and enhancement of selfing in early flowers to ensure reproductive success, are among the most influential traits that contribute to the extremophyte lifestyle of *S. parvula* in its natural habitat.

**One sentence summary:** *Schrenkiella parvula* salt-resilient growth is facilitated by uncompromised primary root growth, expansion of xylem vessels, maintenance of leaf water status and photosynthesis, and early flowering.

## Introduction

Soil characteristics, water availability, and light and temperature regimes set boundaries for plant growth and dictate how plants complete their life cycles. Balancing plant growth with environmental stress responses is a constant challenge faced by all plants. This balance is epitomized by extremophytes, which are plants adapted to thrive in extreme environmental conditions than mesophytes. Therefore, extremophytes provide insights on evolutionarily-tested lifestyle strategies that have proven successful in extreme environments (Lloyd and Oreskes, 2018; Rodell et al., 2018; Schlenker and Auffhammer, 2018; Solis et al., 2020). Knowledge transfer from extremophyte-focused research to design innovative crops that can show resilient growth under stress can be a fruitful strategy when envisioning global food security and expansion of agricultural lands to marginal lands (Oh et al., 2012; Bechtold, 2018; Kazachkova et al., 2018; Zandalinas et al., 2021). The recent genome explorations in a wider range of plants makes this an ideal time for mining representative extremophyte traits.

Salt tolerance is a complex trait that requires integration of physiological, anatomical, and metabolic responses (Cheeseman, 2013; Barros et al., 2021). These multifaceted traits cannot be achieved in stress-sensitive crops simply through the acclamatory response of a few genes (Roy et al., 2014). The complexity of these interactions is implied by the fact that breeding has had limited success in conferring salinity tolerance despite numerous attempts (Yamaguchi and Blumwald, 2005; Cheeseman, 2013; Roy et al., 2014; Barros et al., 2021). An understanding of the extent of tradeoffs between plant growth and stress tolerance is required to assess how selected traits, genes, or pathways affect short- or long-term plant growth and fitness in extremophytes. Such foundational studies are lagging for all the leading extremophyte models compared to the breadth of tools and molecular resources available to identify candidate genes in these species.

*Schrenkiella parvula* (Schrenk) D.A.German & Al-Shehbaz (previously known as *Thellungiella parvula* or *Eutrema parvulum*) is a leading extremophyte model in the Brassicaceae family (Zhu, 2015; Ali and Yun, 2017; Kazachkova et al., 2018; Krämer, 2018). *S. parvula* shares many traits with the model plant, *A. thaliana*, that make it an excellent model, including comparable genome size (Dassanayake et al., 2011), similar lifecycle duration, self-pollination, and prolific seed production (Table 1 and Figure 1). In the ten years following the release of its genome, *S. parvula* has been developed as a model system equipped with primary molecular tools to explore genetic mechanisms underlying plant abiotic stress tolerance (Dassanayake et al., 2011; Oh et al., 2014; Wang et al., 2018; Pantha et al., 2021; Wang et al., 2021). *Schrenkiella parvula* is remarkably tolerant to high salinity compared to closely related Brassicaceae species (Orsini et al., 2010) and is categorized as a halophyte based on its capacity to complete its life cycle when grown at 200 mM NaCl or higher (Flowers and Colmer, 2008). In addition to high sodium, *S. parvula* is also uniquely adapted to cope with multiple edaphic factors found in its native habitat, including boron, potassium, and lithium, among other salts, that exist at levels toxic for most crops (Helvaci et al., 2004; Oh et al., 2014; Hajiboland et al., 2018; Tug et al., 2019). Its preferential distribution near saline lakes in the Irano-Turanian region makes it a robust extremophyte model to investigate multiple environmental stresses that are often found as compound stresses in many marginal agricultural landscapes.

**Figure 1.**
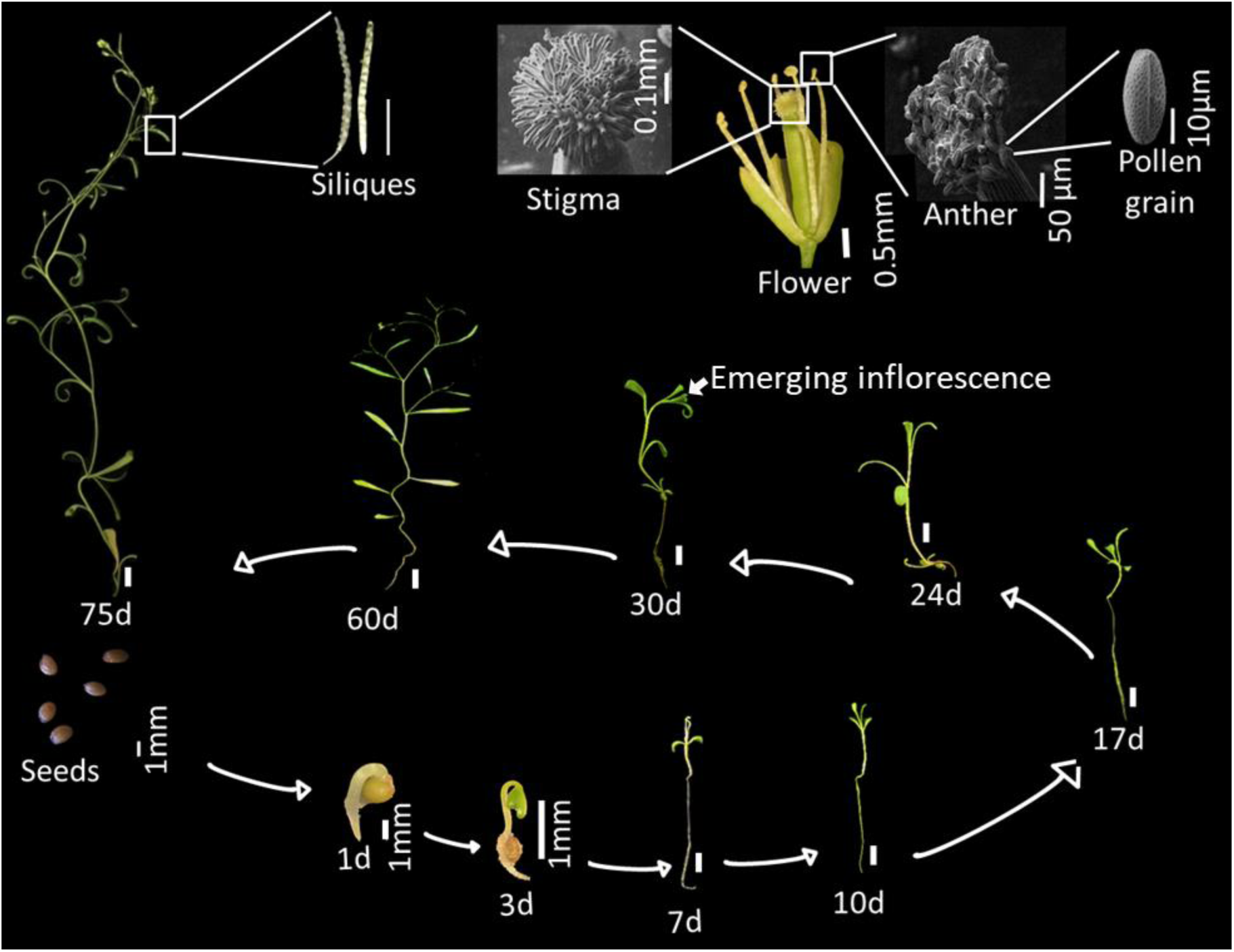
Life cycle of *Schrenkiella parvula* from seeds to siliques. Scale bars are 1 cm unless indicated in the figure.

**Table 1.**
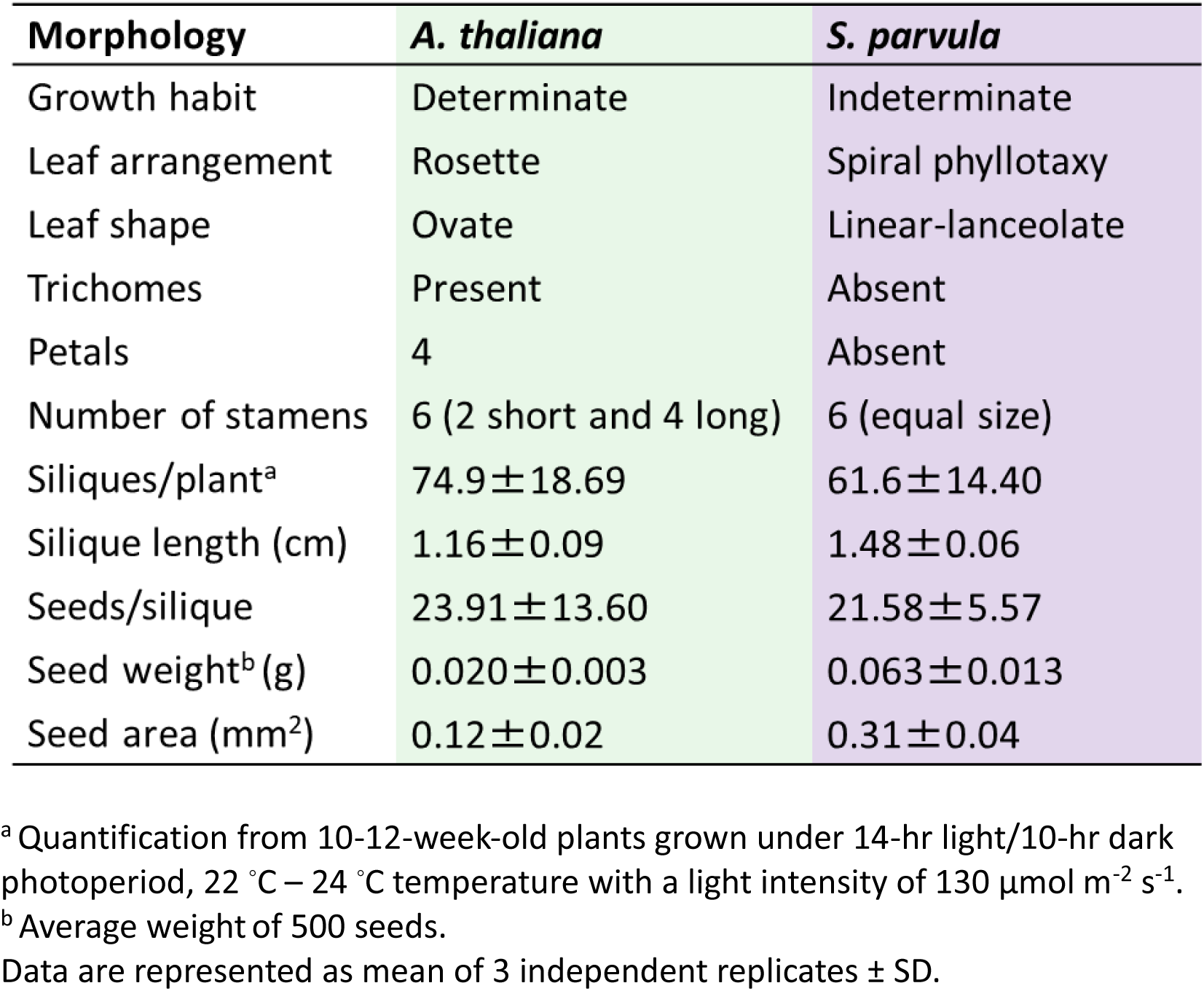
Morphological comparison between *Arabidopsis thaliana* and *Schrenkiella parvula*.

Despite a growing body of genomic resources for *S. parvula* (Jarvis et al., 2014; Ali et al., 2018; Wang et al., 2021), a systematic study that describes its life history, and the physiological, and structural features that are associated with adaptations to environmental stresses, especially salt stress, is absent. In this study, we evaluated *S. parvula* stress adaptations at both the structural and physiological levels. We assessed adaptations that are likely to be influential in balancing growth with stress tolerance and identified inducible adaptive traits that were the most prominent features in a multivariate space across developmental stages, tissue types, and response types. We propose these inducible traits to be hallmarks of a stress-adapted lifestyle for a fast-growing annual in saline soils demonstrating the utility of *S. parvula* as an extremophyte model plant.

## Results

### *Schrenkiella parvula* adjusts root growth, structure, and form under high salinity

*Schrenkiella parvula* roots, in contrast to those of *A. thaliana*, maintained uninterrupted root growth in response to salt stress compared to control conditions (Figure 2A). Remarkably, *S. parvula* primary root length did not display any growth retardation even after longer durations or higher salt concentrations tested, while *A. thaliana* showed growth inhibition at 100 mM NaCl (Figure 2B). The same trend was observed for lateral root growth, although the emergence of lateral roots was delayed in *S. parvula* even at control conditions compared to *A. thaliana* (Figure 2C). The higher tolerance to salinity shown by *S. parvula* roots compared to *A. thaliana* roots was consistent when plants were grown under a longer photoperiod and at higher nutrient levels (Figure S1). Under these conditions, a 125 mM NaCl treatment caused a growth stimulating effect on *S. parvula* primary root length, but inhibited root growth in *A. thaliana* (Figure S1B). Stronger inhibition of both average lateral root length and number was imposed by 175 and 225 mM NaCl, which could explain the reduced total lateral root length (Figure S1C to E). Taken together, salt stress imposed by 125 mM NaCl promoted primary root growth and average lateral root length, while a higher salt concentration of 175 mM NaCl imposed less inhibition on main root growth and lateral root density in *S. parvula* compared to *A. thaliana*. Compared to primary and lateral roots, root hair development in *S. parvula* was sensitive to high salinity; root hairs showed a consistent decrease in length at higher salinities or during longer exposure times to salt stress (Figures 2D and S2). The ability of *S. parvula* to maintain or enhance primary root growth under high salinity remained as a consistent trait shown by mature plants over longer durations and higher salinities tested using 250 mM NaCl in a hydroponic growth medium (Figure S3).

**Figure 2.**
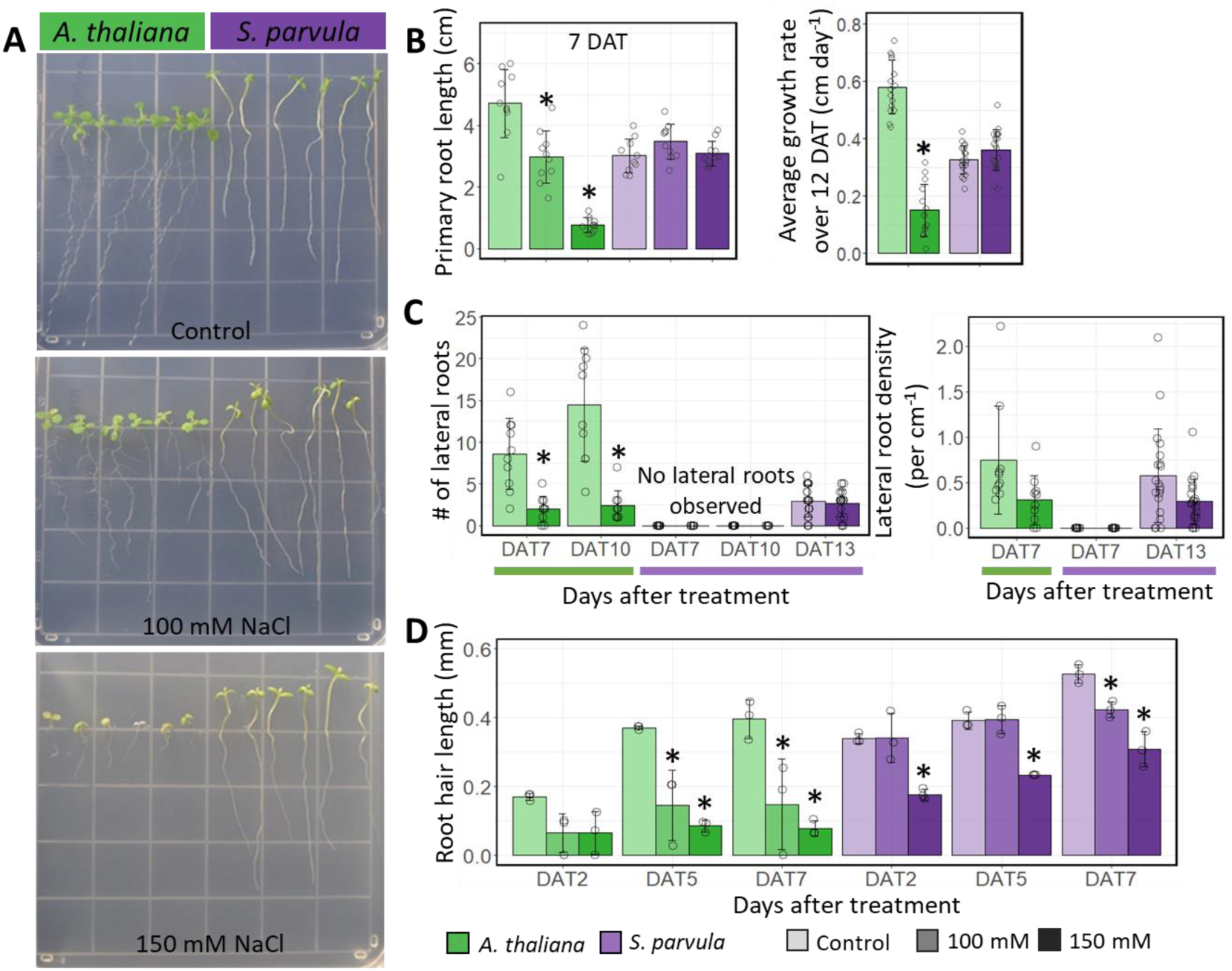
Effects of NaCl stress on root growth in *Schrenkiella parvula* and *Arabidopsis thaliana* seedlings. [A] 12-day-old seedlings of *A. thaliana* and *S. parvula* were grown for 5 days on 1/4x Murashige and Skoog media, and 7 days on the indicated concentrations of NaCl. Plates were scanned 7 days after treatment (DAT). [B] Primary root growth and average root growth rate, [C] number of lateral roots and lateral root density, and [D] average length of 10 longest root hairs. Asterisks indicate significant difference (*p* ≤ 0.05) between the treated samples and their respective control group, determined by Student’s *t*-test. Data are mean ± SD (n =3, at least 3 plants per replicate). Open circles are individual measurements.

*Arabidopsis thaliana* roots showed halotropism by growing away from salt (Figure S4) (Galvan-Ampudia et al., 2013). Interestingly, under comparable growth conditions, *S. parvula* showed salt-insensitive primary root growth where it grew towards higher salt concentration without changing its course, further exemplifying the higher tolerance to high salinity (Figure S4).

To investigate the potential effect of salt stress on root anatomy, we examined the tissue level structural responses in roots of 8-week-old *S. parvula* plants that had been treated for 4 weeks with 150 mM NaCl (Figure 3A). These traits were catalogued from root tips to mature roots (Figure S5). In young roots, the length of the root tip (measured as the distance from the tip to the emergence site of the first root hair) was longer under high salinity conditions compared to control conditions (Figure 3B). However, we did not observe any other anatomical trait adjustments to salinity in young roots of *S. parvula* (Figure 3C). Interestingly, *S. parvula* roots develop an extra cell layer between the cortex and endodermis compared to *A. thaliana* (Brady et al., 2007), formed early on in its root development in both control and salt-treated samples (Figure S5A and C). In response to high salinity, xylem tissues in mature roots significantly increased in area under high salinity (Figure 3D). However, this did not affect the overall mature root area. The xylem tissue expansion was primarily caused by increase in the area of vessels. The extra space taken up by the xylem tissue was compensated for by reduction of the cortical air spaces (unstructured aerenchyma) in mature roots (Figure 3D).

**Figure 3.**
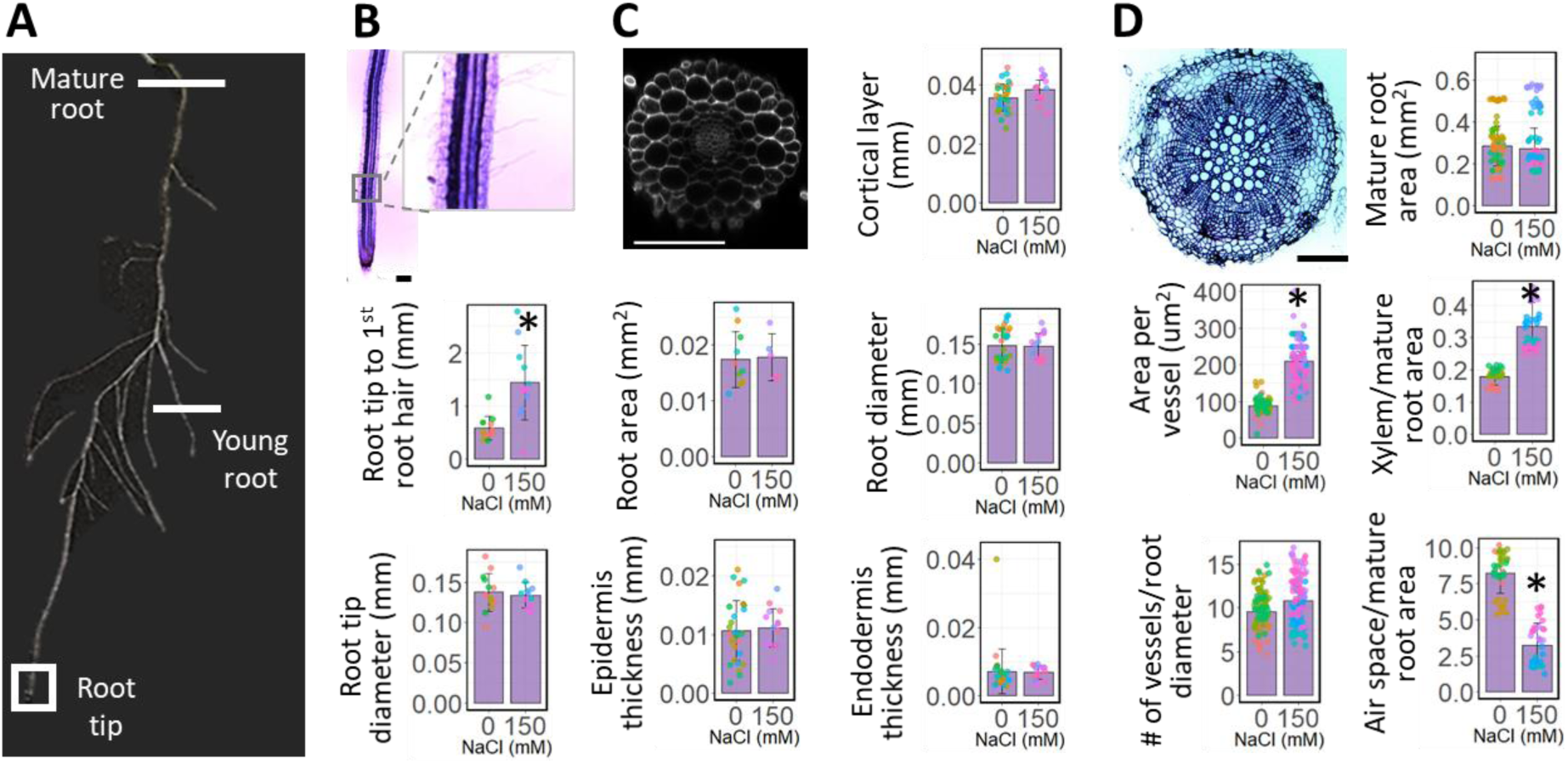
Effects of NaCl stress on *Schrenkiella parvula* root anatomy. Eight-week-old *S. parvula* plants were grown under control or 150 mM NaCl conditions for an additional 4 weeks. [A] The position of transverse sections of *S. parvula* roots, [B] the root tip, [C] the young root, and [D] the mature root from 12-week-old hydroponically-grown *S. parvula* plants. A minimum of 20 sections from 4 to 13 plants were examined for each root region to quantify each parameter for each condition. Asterisks indicate significant differences (*p* ≤ 0.05) between the treated samples and their respective control group, determined by Student’s *t*-test. Data are means ± SD. Data points represent individual cross-sections and colors represent individual plants. Representative cross-sections were obtained from the control plants. Scale bar represents 100 µm.

### *Schrenkiella parvula* shoots change in form and function to response to salt stress

Structural adjustments in response to long-term salt stress extended to shoot tissues in *S. parvula* (Figures 4 and S6). Long-term salt stress leads to larger vessels in *S. parvula* shoots (Figure 4A) to maintain the root-shoot continuum with root vessel elements that increased in size when exposed to high salinity (Figure 3D). The expansion in vessel element size in shoots was matched by a reduction in cambial and cortical zones that kept the average shoot area constant between control and salt-treated plants (Figure 4A), while other cell and tissue layers in the shoot remained unchanged (Figure S6B).

**Figure 4.**
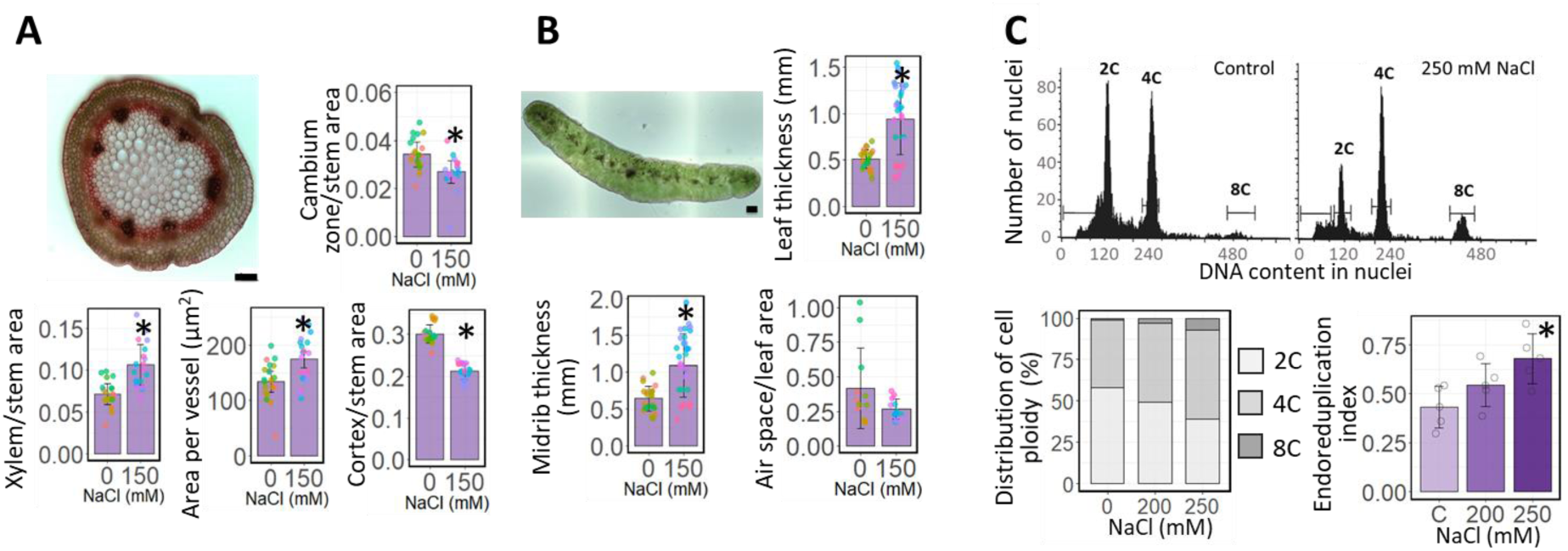
Effects of NaCl stress on *Schrenkiella parvula* shoot anatomy. Eight-week-old *S. parvula* plants were grown under control or 150 mM NaCl conditions for an additional 4 weeks. Transverse section of [A] stem and [B] leaf of 12-week-old hydroponically-grown *S. parvula* plants. A total of 23 sections from 7 control plants and 32 sections from 9 treated plants were used for leaf measurements, and a minimum of 20 sections from 5 to 11 plants were used for stem measurements. Data points represent individual cross-sections and colors represent individual plants. [C] Nuclear DNA content, distribution of leaf cell ploidy, and endoreduplication index of leaf cells from 10^th^ and 11^th^ leaves from the shoot meristem in control and 250 mM NaCl-treated 8-week-old *S. parvula* plants. Data are mean ± SD (n= 4). Asterisks indicate significant difference (*p* ≤ 0.05) between the treated samples and their respective control group, determined by Student’s *t*-test. Representative cross-sections were obtained from the control plants Scale bars represent 100 µm.

Leaves that developed during long-term salt treatments in *S. parvula* exhibited increased succulence, as indicated by increase in leaf thickness across the leaf including the midrib (Figures 4B and S7). Increased endoreplication induced by salinity can cause cell size expansion and succulence in the extremophyte, *Mesembryanthemum crystallinum* (Barkla et al., 2018). We therefore tested whether the increased succulence in *S. parvula* leaves in response to salt coincided with endoreplication by quantifying ploidy of leaf cells. Indeed, the proportion of leaf cells with higher ploidy increased with increased salinity, causing a significant rise in the endoreplication index under 250 mM NaCl compared to the control condition (Figure 4C). *Schrenkiella parvula* leaves showed both structural and functional adjustments to salt treatments that correlated with its capacity to maintain relative water content (Figure 5), but the total leaf number per plant remained similar between control and salt-treated plants (Figure 5A). Long-term salt treatments not only increased leaf succulence, but also leaf area (Figure 5A), suggesting a growth-promoting effect at least with the long-term 150 mM NaCl treatment. *Schrenkiella parvula* was photosynthetically active and maintained growth during long-term salt treatments at high salinities (Figure 4), but there was a tradeoff in increasing salt on shoot development at salinities surpassing 150 mM (Figure S8).

**Figure 5.**
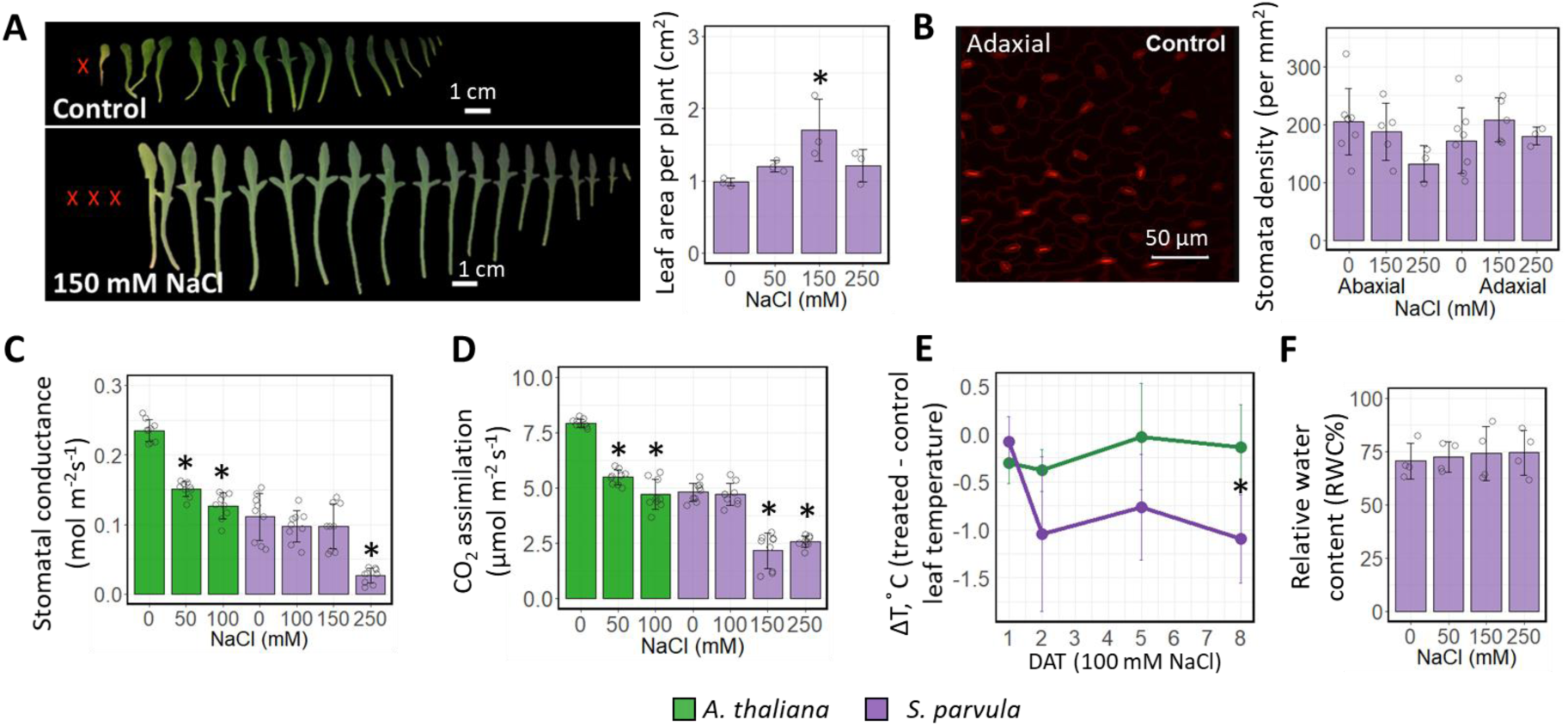
Effects of NaCl stress on *Schrenkiella parvula* leaf traits. All experiments were performed with 4-week-old hydroponically-grown plants that were treated for an additional 4 weeks with the indicated NaCl concentrations. [A] Total leaf area, [B] Stomatal density, [C] Stomatal conductance, [D] Photosynthesis rate, [E] Leaf relative surface temperature, and [F] Leaf relative water content. Asterisks indicate significant difference (*p* ≤ 0.05) between the treated samples and their respective control samples, determined by Student’s *t*-test. Data are means ± SD (n ≥ 3). Open circles indicate the measurement from each plant (in B and F) and leaves (in C and D) for each experiment. DAT, days after treatment.

Leaf shape, angle and phyllotaxis, stomatal distribution, and boundary layer thickness caused by thick cuticles or trichomes largely contribute to efficient use of water by plants (Yoo et al., 2009). *Schrenkiella parvula* leaves are amphistomatous (i.e. stomata exist on both abaxial and adaxial surfaces), linear-lanceolate with a narrower base, and trichomeless (Table 1, Figure 5A and 5B). In *S. parvula*, stomatal density did not change in fully-mature leaves (Figure 5B). However, stomatal regulation was adjusted with increasing salinities as indicated by decreased stomatal conductance (Figure 5C) and decreased CO_2_ assimilation, notably at much higher salinities for *S. parvula* than observed for *A. thaliana* (Figure 5D). While 50 mM salt treatment was sufficient to cause a decrease in stomatal conductance and photosynthesis in *A. thaliana*, *S. parvula* treated with 100 mM NaCl was able to maintain stomatal conductance and CO_2_ assimilation at levels indistinguishable from those measured in plants grown under control conditions. Concomitantly, *S. parvula* maintained lower average leaf temperatures than *A. thaliana* (Figure 5E). Remarkably, *S. parvula* maintained relative leaf water content at all tested salinities up to 250 mM NaCl (Figure 5F).

Utilizing the Qubit/PSI PlantScreen^TM^ Compact high-throughput phenomics system, we undertook a comparative analysis of *A. thaliana* and *S. parvula* shoot morphometrics (RGB digital camera), leaf water content (infra-red camera), and photochemistry (PAM chlorophyll fluorescence). Figure S9A demonstrates that the phenomics system was able to simultaneously capture morphological characteristics specific to each species. *A. thaliana* has a rosette structure (i.e. less protrusions from a perfect circle), and therefore exhibited higher compactness). In contrast *S. parvula* leaves on a spiral phyllotaxy (Figure 1 and Table 1) that increasingly protrude as the plant grows, reflected a lower compactness in older plants regardless of salt concentrations. Salt stress led to a statistically significant reduction in *A. thaliana* leaf area but had no effect on *S. parvula* leaves (Figure S9B). *S. parvula* was also able to maintain leaf water content and maximum quantum efficiency of PSII under salt stress (Figure S9C and D, respectively) whereas *A. thaliana* exhibited a salt stress-mediated reduction in both parameters. Non-Photochemical Quenching (NPQ), a mechanism for dissipating excess light energy (Müller et al., 2001), increased in *A. thaliana* under salt stress suggesting that less energy is being utilized for productive purposes (Figure S9E). On the other hand, salt stress had no effect on NPQ in *S. parvula*. The resilience of *S. parvula* photochemistry under salt stress is consistent with this species’ ability to maintain CO_2_ assimilation in saline conditions (Figure 5D). Taken together, the phenomics data confirm similar growth and physiological tolerance of *S. parvula* to salt stress regardless of growth medium.

Minimizing non-stomatal transpiration by enhanced epidermal boundary layer resistance is equally important as stomatal regulation in preventing water loss under water stress (Blum, 2009). Indeed, previous findings have demonstrated that *S. parvula* leaves possess a significantly thicker leaf cuticle supported by a higher level of total wax content compared to *A. thaliana* leaves (Teusink et al., 2002) (Figure S10A). Interestingly, when we compared the basal expression of key genes in the wax biosynthesis pathway (Bernard and Joubès, 2013) from comparable shoot tissues grown under control conditions, we observed consistently higher constitutive expression in *S. parvula* compared to expression of their *A. thaliana* orthologs (Figure S10B).

Changes to leaf shape and arrangement is not a salt-dependent, inducible trait in *S. parvula*. However, its narrow leaves with elongated petioles arranged spirally on erect stems with elongated internodes (Figure S11A) are ideally suited to rapid growth in saline habitats with warm temperatures where efficient transpirational cooling imparts selective advantages (Lin et al., 2017). Additionally, an erect growth habit is more desirable than a rosette to minimize leaf contact with soil, given that *S. parvula* is found near saline lakes with topsoils often enriched in salt crystals (Hajiboland et al., 2018; Tug et al., 2019). Key traits resulting in narrow leaves, elongated petioles and internodes are influenced by the phytochrome family of genes (Somers et al., 1991; Li et al., 2011), and we noted that *S. parvula* plants closely resemble the morphology of *A. thaliana phyB phyD* double mutants (Devlin et al., 1996; Li et al., 2011). This morphological similarity led us to search for genome-level cues for loss-of-function or altered function in *S. parvula PHY* genes that may support such a phenotype being selected as the only documented growth form of *S. parvula* (Figure 1 and Table 1) (German and Al-Shehbaz, 2010). We observed that *PHYB* was conserved as a single copy gene in six closely-related Brassicaceae genomes that we examined, with a translocation event between lineage I and II species. However, *PHYD* appeared to have undergone a gene loss specifically in *S. parvula* (Figure S11B).

### Salt stress induces early flowering and silique formation in *Schrenkiella parvula*

We examined long-term salt-stress effects on reproductive traits to investigate how excess Na^+^ affected *S. parvula* fitness (Figure 6). Salt treatment induced early flowering irrespective of whether flowering time was measured as the number of days after planting to detection of the first floral buds or the number of leaves developed before flowering (Figure 6A to C). Additionally, early flowering induced by salt was observed under both long-day and short-day photoperiods (Figure 6C). The initial ∼10 flowers produced under control conditions aborted without developing into siliques, while these flowers produced mature siliques in salt-treated plants (Figure 6A and B). *Schrenkiella parvula* flowers are self-fertilizing, a process facilitated by the elongation of filaments to the level of the stigma. However, early flowers under control conditions developed shorter filaments than in the salt-treated plants (Figure 6D). In control plants, these shorter filaments appeared to hinder successful fertilization and delayed subsequent silique formation (Figure 6A and E). Notably, the salt-induced early flowering and salt-dependent elongation of filaments were not specific to NaCl treatment and were also initiated by KCl at similar concentrations (Figure S12). At later reproductive stages, the continuous growth of multiple floral meristems led to fertile flowers under control conditions. Despite the early success of silique formation in salt-treated plants, the cumulative number of flowers and siliques produced by control plants surpassed that of salt-treated plants if allowed to grow past two months. For example, at 74 days after planting, the control plants had a greater number of flowers per plant than salt-treated plants (Figure 6E). The experiment was discontinued when salt-treated plants stopped producing new inflorescences and were senescing (approximately 74 days after planting). Therefore, *S. parvula* does not require salt to increase fitness, and salt treatments appear to only provide a transient advantage during early reproductive stages. Even if fitness is higher for control plants compared to salt-treated plants when evaluated based solely on the total number of siliques produced per plant within a longest possible growing season, there may be additional benefits when seeds develop on plants exposed to long-term high salinity, especially if seasonal changes in its natural habitats reward a strategy of salt-accelerated reproduction. The average size of siliques and seeds of salt-treated plants were significantly greater than in control plants (Figure 6G, H, and I). This suggested that more resources were available for seeds when produced by salt-treated plants compared to control plants. The number of seeds per silique was not different between control and treated plants and the seeds from both groups were indistinguishable in the rate of germination in the subsequent generation when germinated on media without added NaCl.

**Figure 6.**
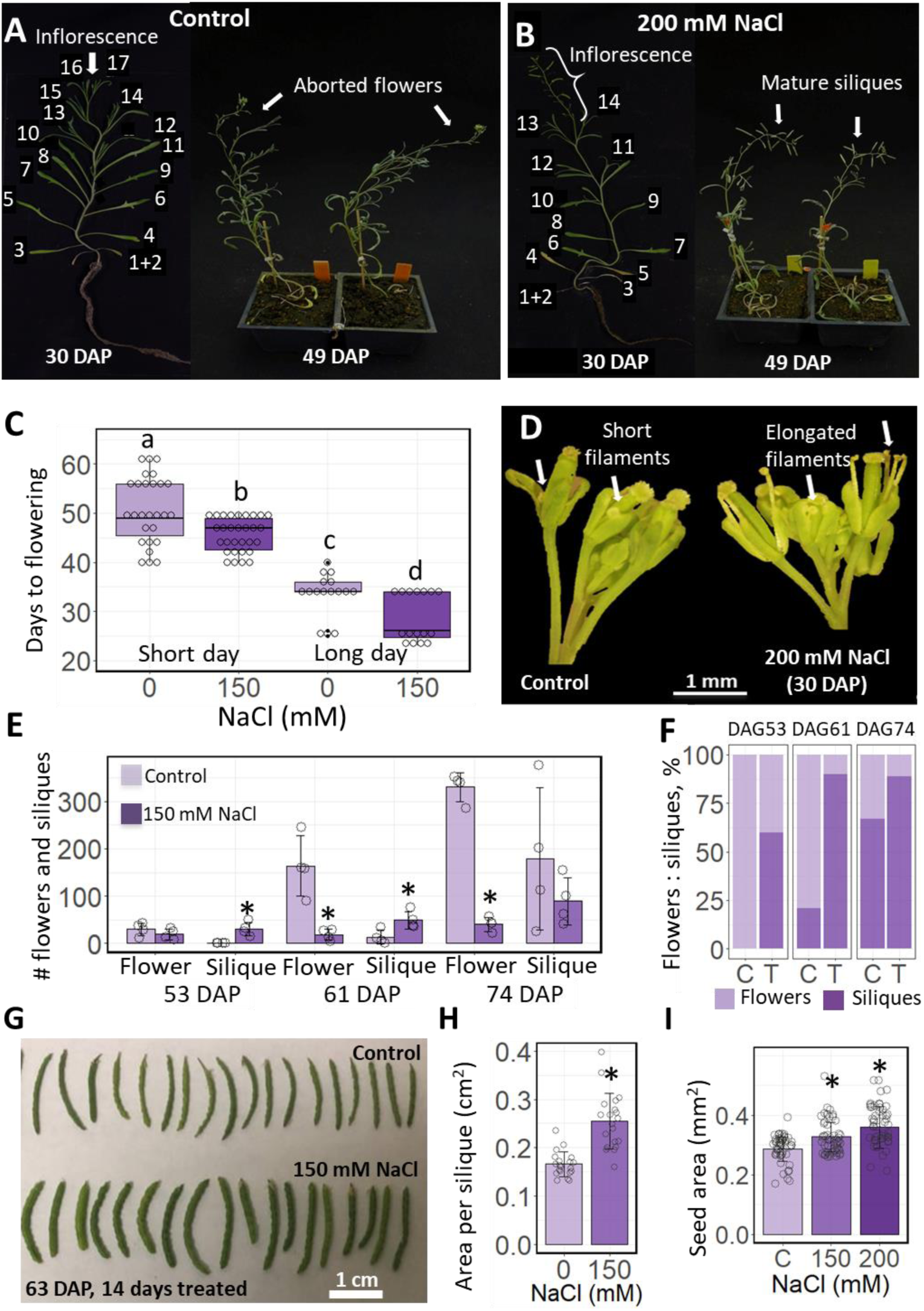
Effects of NaCl on reproductive traits of *Schrenkiella parvula*. [A] Plants grown under control conditions generally flower at the ∼17th leaf stage and the first few flowers are subsequently aborted. [B] Salt treated plants flower earlier at the ∼14th leaf stage and the first flowers develop into mature siliques. [C] Days from planting to the first observed flower of plants grown under a long day (16 hr light/8 hr dark) or short day (12 hr light/12 hr dark) photoperiod with and without the indicated salt treatment. Center line - median; box - interquartile range (IQR); notch - 1.58 × IQR/sqrt(n); whiskers - 1.5 × IQR. [D] Flowers obtained from control and treated plants described in panel A and B. [E] Number of flowers and siliques per plant under control and salt treatment. Data are mean ± SD. Open circles represent individual measurements from four biological replicates. [F] Ratio between the number of flowers and siliques observed for control (C) and salt-treated (T) plants described in E. [G] Siliques from plants under control and NaCl treated conditions. [H] Size of siliques from G. Open circles represent individual silique counts. [I] Comparison of the area of seeds from plants under control and NaCl treated conditions. Data are mean ± SD (n = 50). Seeds per condition were selected from the seed pool of 15 to 50 plants. Open circles represent individual seeds. For panel [B], salt treatment started 21 DAP, initially at 50 mM NaCl given every other day. The salt concentration was increased by 50 mM every four days until it reached 200 mM; for panels [C, E, F], salt treatment was applied to 3-week-old plants until the end of the experiment; for panels [G, H, I], salt treatment was applied to 5-week-old plants for an additional 2 weeks. Different letters or asterisks represent significant differences (*, *p* < 0.05) compared to control, determined by either one-way ANOVA followed by Tukey’s post-hoc test [C] or Student’s *t*-test [E, H, I]. DAP, days after planting.

### *Schrenkiella parvula* seed germination is delayed by high salinity and is sensitive to specific salts

Many plants including salt-adapted plants delay germination in saline media even if their seedling stages can tolerate high salinity (Ungar, 1978; Kazachkova et al., 2016). Both *S. parvula* and *A. thaliana* exhibited decreased seed germination as NaCl concentrations increased compared to control conditions (Figure 7A). However, unlike the more salt-tolerant traits *S. parvula* displayed compared to *A. thaliana* during seedling and mature developmental stages (Figures 2, 4, and S9), *S. parvula* seed germination was more sensitive to NaCl than *A. thaliana* seed germination (Figure 7A). When we sowed *S. parvula* seeds directly onto salt plates prior to stratification, we observed more severe salt-mediated inhibition of germination. This was demonstrated via detailed germination curves that revealed a progressive reduction in both germination rate and final percent germination with increasing NaCl concentrations (Figure 7B).

**Figure 7.**
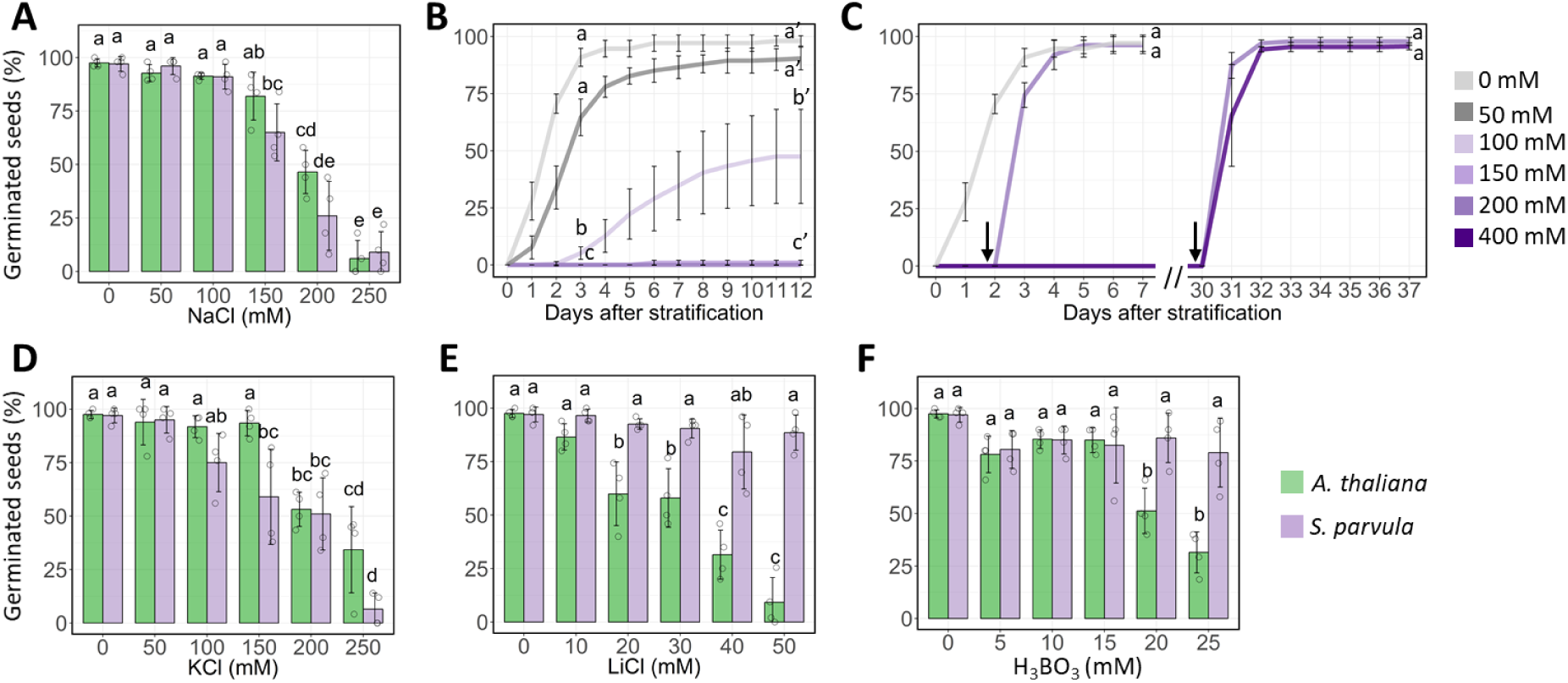
Effects of salt stress on the germination of *Arabidopsis thaliana* and *Schrenkiella parvula* seeds. [A] Final percent germination recorded at 3 days and [B] germination curves of seeds treated with different concentrations of NaCl recorded for 12 days after stratification. [C] Germination curves of NaCl-treated seeds transferred to control (no NaCl) media, after 2 days or 30 days on salt plates. The germination curves were performed by direct sowing of seeds onto salt plates prior to stratification. Black arrows point to the day when seeds were transferred to non-NaCl plates. [D, E, F] Final percent germination of seeds treated with KCl [D], LiCl [E] or H_3_BO_3_ [F]. Data are mean ± SD (n = 3-4, each replicate contains ca. 50 seeds). Open circles indicate individual measurements. Different letters indicate significant difference (*p* ≤ 0.05) determined by one-way ANOVA with post-hoc Tukey’s test across all conditions in [A, D, E, F], or across treatment groups at 3 and 12, and 7 days after stratification in [B] and [C] respectively.

This salt-influenced trait did not lead to a binary outcome between germinated vs ungerminated seeds. The smaller fraction of *S. parvula* seeds that formed a radicle compared to *A. thaliana* continued to develop in a high salinity medium similarly to those seeds germinated under control conditions (Figure S13A). In contrast, *A. thaliana* seeds that formed radicles in a high saline medium did not survive to the seedling stage. Up to 98% of *S. parvula* seeds that failed to germinate (i.e. no visible radicle emergence) on high-salt growth medium germinated and developed into seedlings when transferred to control medium without added NaCl (Figures 7C and S13B). Moreover, seeds that were incubated for a month on saline medium still exhibited 98% germination (Figure 7C). Interestingly, adding salt and increasing the duration of salt-treatment, led to faster germination when salt-inhibited seeds were transferred back to the control medium. These results suggest that NaCl introduces a reversible barrier to *S. parvula* seed germination.

We also tested whether other salts, including K, Li, and B, found in the native soils of *S. parvula* ecotype Lake Tuz caused a reversible inhibition of *S. parvula* seed germination. KCl induced a similar inhibitory response as observed for NaCl (Figures 7D and S13), but neither LiCl nor H_3_BO_3_ led to inhibition of *S. parvula* seed germination (Figure 7E and F). In contrast, *A. thaliana* seed germination was highly sensitive to high LiCl and H_3_BO_3_, and KCl at 150 or 250 mM was lethal to *A. thaliana* seeds that formed a radicle (Figures 7D, E, F, and S13A). These data suggest that *S. parvula* is adapted to sense the soil salt level as well as the salt type before radicle emergence, and if seeds germinate, they are more likely to continue development despite toxic levels of salts in the growth medium that are lethal to *A. thaliana*.

Salt-dependent germination of *S. parvula* suggests that it is an adaptive trait to time seed germination for when topsoil salinity decreases during the summer months from June to September. This finding was supported by field observations made in the Lake Tuz region where *S. parvula* grows as a seasonal annual during the summer months that receive the highest rainfall and temperature with the longest day length (Figures S14A to C). The saline soils surrounding Lake Tuz are often covered with a salt crust towards fall, which becomes a dominant feature in the landscape extending to spring of the following year despite having a high saline water table throughout the year (Tug et al., 2019). Seed germination may range widely within the expected growing season and seedlings are more vulnerable to extreme environmental conditions than mature plants. Finally, we compared *S. parvula* and *A. thaliana* seedlings for their tolerance to heat, chilling, and freezing stresses to assess how *S. parvula* may fit within the expected temperature tolerance range required to survive in its native habitat (Figures S14D to F). *Schrenkiella parvula* seedlings displayed higher tolerance to heat stress at 38 °C than *A. thaliana* (Figure S14F), while chilling and freezing tolerance in both species was indistinguishable (Figure S14D and E).

## Discussion

Plants are known for their remarkable capacity to tolerate environmental stresses and ability to modulate growth even to the extent of pausing growth entirely. This plasticity in response to various environmental stresses is observed in all plants, but as shown in the current study, extremophytes such as *S. parvula* exhibit a much higher tolerance to stress than mesophytes, which include most crops and *A. thaliana* (Flowers and Colmer, 2008; Kazachkova et al., 2018). Halophytes represent only about 0.4% of all flowering plants and about 40% of all halophytes including *S. parvula* can withstand salt stress at concentrations similar to seawater (Kotula et al., 2020). At high salinities, most plants respond to salt stress by inhibiting growth to prioritize survival as a tradeoff (Santiago-Rosario et al., 2021). Nevertheless, the extremophyte model *S. parvula* provides a genetic system to discover adaptive traits that do not show a growth compromise at salt concentrations that are known to be lethal to most crops and *A. thaliana* (Oh et al., 2014). The defining traits of *S. parvula* as an extremophyte should reflect the environmental constraints that have shaped its phenotype and collectively made it distinct from other more stress-sensitive annuals in Brassicaceae, including *A. thaliana*. When we summarized *S. parvula* physiological and structural traits that changed at a developmental stage-specific and tissue-specific scale in response to salt stress, several traits emerged as significant features induced under salt stress compared to their non-stressed control group (Figure 8A). Among them, the higher number of siliques produced during salt-induced early flowering, expansion of xylem vessel elements, and increased leaf thickness provide the greatest contribution to the total plastic or inducible trait space (Figure 8B and C).

**Figure 8.**
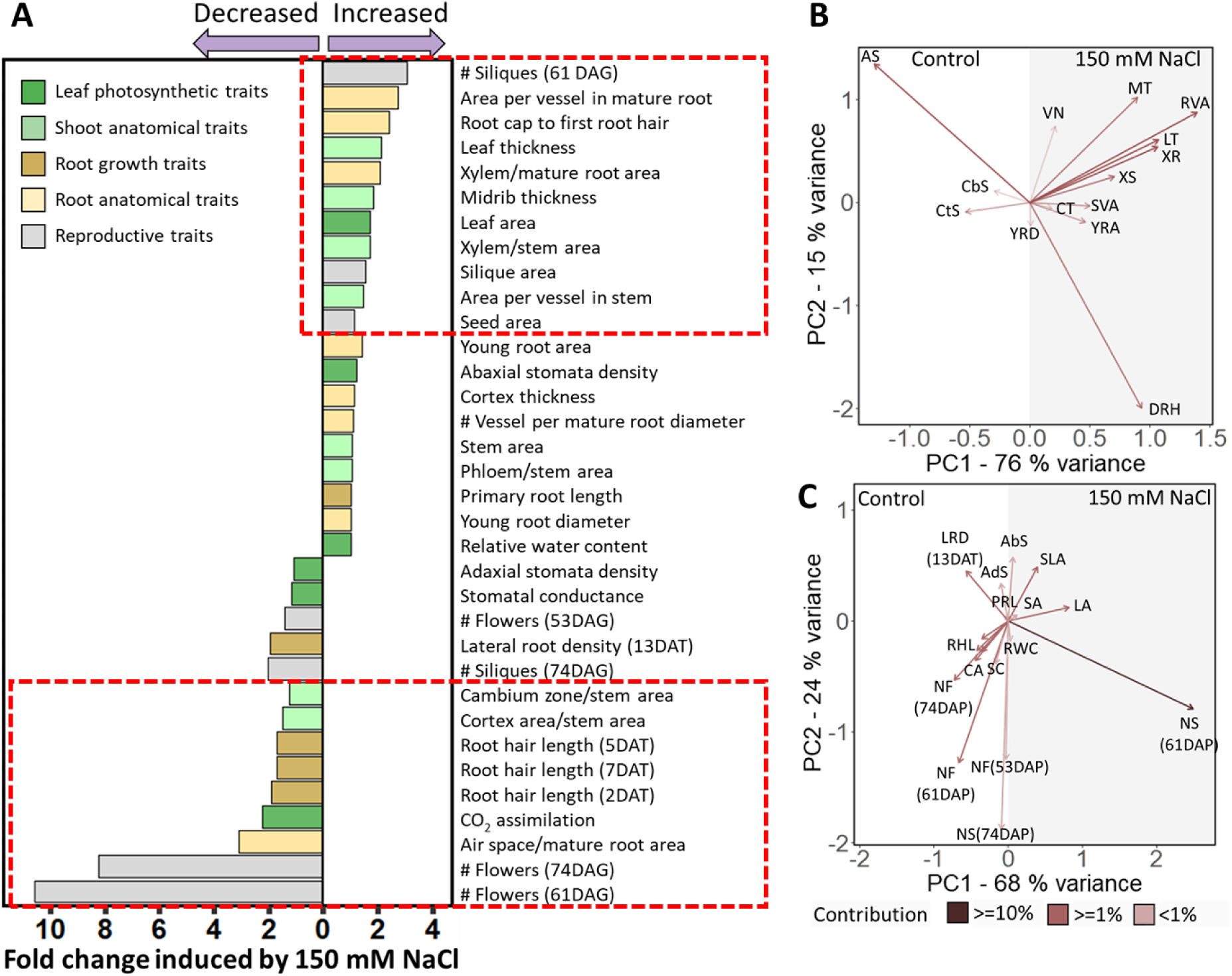
Summary of salt-induced structural and physiological traits in *Schrenkiella parvula*. [**A**] Salt-induced changes in traits quantified in the current study. Changes in each trait were calculated as the fold change between treatment and control measurements. Traits highlighted in the red boxes showed significant differences (*p* < 0.05) under salt treatments compared to the control determined in previous assays (Fig 2-7). [**B**] and [**C**] PCA biplot of traits quantified for *S. parvula* under control and salt-treated conditions. Arrows indicate directions of loadings for each trait and are color-coded by contribution to the variations in PC1 and PC2. [B] Anatomical traits quantified: LT, leaf thickness; MT, midrib thickness; XS, stem xylem to stem area; SVA, average area per vessel in stems; CbS, cambium to stem area; CtS, cortex to stem area; XR, root xylem to root area; RVA, average area per vessel in roots; VN, number of vessels across the root diameter; AS, air space to root area; YRA, young root area; CT, cortical thickness; YRD, root diameter in young roots; DRH, distance from root tip to first root hair. [C] Physiological traits quantified: AbS, abaxial stomatal density; adS, Adaxial stomatal density; LA, leaf area; SLA, silique area; SA, seed area; PRL, primary root length; LRD, lateral root density; RWC, relative water content; CA, CO2 assimilation; SC, stomatal conductance; RHL, root hair length; NF, number of flowers; NS, number of siliques. DAT, days after treatment; DAP, days after planting.

### Salt stress accelerates reproduction thereby contributing to maximizing fitness of *S. parvula*

Salinity delays flowering in most plants (Blits and Gallagher, 1991; Lutts et al., 1995; Apse et al., 1999; Moriuchi et al., 2016; Cho et al., 2017). For example, high salinity (≥100 mM NaCl) delays or inhibits the transition from vegetative to reproductive growth in *A. thaliana* (Achard et al., 2006; Li et al., 2007; Ryu et al., 2014). Delayed flowering ensures survival under salt stress supported by multiple genetic mechanisms. These pathways include transcription factors that inhibit flowering and are also induced by salt (Cho et al., 2017). However, a few plants possess alternative strategies whereby flowering is accelerated in response to salt stress (Adams et al., 1998; Ventura et al., 2014). Additionally, molecular pathways operating antagonistically to salt-induced delayed flowering have been described for *A. thaliana* (Yu et al., 2018). This apparent strategy would facilitate reproduction and improve fitness by allowing escape from harsh environments, especially for ruderal or annual plants growing in habitats that become warmer, drier, and more saline (due to increased surface evaporation) towards the end of their growing season. Salt-induced flowering observed for *S. parvula* suggests that such an alternative strategy was selected as the dominant trait to maximize its fitness in its native habitat exemplified by the Lake Tuz ecotype used in our current study (Figures 6 and S12) over the more common trait of salt-induced delay in flowering observed for many other plants (Apse et al., 1999; Cho et al., 2017).

Compared to favorable growth conditions, environmental stresses increase the opportunity for selection. Previous studies have demonstrated that phenotypic selection favors stress-avoidance traits, including earlier flowering, in addition to stress-tolerance traits (Eshel et al., 2021). This results in an evolutionary shift toward earlier flowering despite the prevalent trait observed for model and crop plants to delay flowering when exposed to salt (Stanton et al., 2000; Ventura et al., 2014; Caño et al., 2016; Moriuchi et al., 2016). Stanton et al., (2000) further showed that, among multiple environmental stresses tested for their effects on evolutionary selection in wild mustard, *Sinapis arvensis* (a ruderal annual in Brassicaceae), the potential for selection was greatest for high salt stress and low light. *Schrenkiella parvula* is native to the variable shoreline habitats of saline lakes of the Irano-Turanian region where it is often a ruderal plant growing in frequently inundated saline soils (Ozfidan-Konakci et al., 2015; Hajiboland et al., 2018; Tug et al., 2019). Therefore, salt stress is the norm for *S. parvula* and over a multi-generational time scale, salt stress is expected to provide a constant selection pressure.

When we examined salt-responsive traits throughout the lifecycle of *S. parvula*, an increased number of siliques produced due to salt-induced early flowering had the largest fold-change among other salt-induced traits that differentiated the response between control and salt-treated growth (Figure 8C). Early flowering established an earlier seed set and the opportunity to produce viable seeds at an earlier developmental age in the salt treated plants compared to control plants (Figure 6). However, this was a plastic trait not necessarily imparting higher fitness when assessed based on overall fecundity during a longer growth period. When control and salt-treated plants were grown for over 2 months, the control plants displayed longer vegetative growth phases subsequently leading to higher fecundity in *S. parvula.* These results suggest that salt-induced flowering has evolved as a mechanism of stress avoidance rather than stress tolerance in a habitat where the severity of environmental stresses increase as the growing season continues (Figure S14).

A key morphological feature contributing to the transient advantage of salt-mediated early flowering in *S. parvula,* was the plastic response observed for filament elongation that was enhanced by salt (Figures 6D and S12). Filament elongation is an important trait for successful fertilization in self-pollinating plants such as *S. parvula* and *A. thaliana*. However, whereas salt stress terminally inhibits *A. thaliana* gamete formation and leads to abortion of flowers and seeds (Sun et al., 2005; Figure 6), filament elongation and production of viable seeds were not inhibited by salt in *S. parvula*. This finding indicates that *S. parvula* has evolved to decouple the regulatory pathways that sense salt and inhibit stamen growth.

### Salt stress-induced xylem vessel expansion across the root-shoot continuum could contribute to the salt resilient growth of *Schrenkiella parvula*

Selection on vessel diameter favors narrower vessels to minimize the risk of cavitation independently of ancestry or habitat for a wide range of angiosperms from rainforests to deserts (Olson and Rosell, 2013). Furthermore, the effect of salt stress on xylem development within a species often leads to narrower vessels (Junghans et al., 2006; Ge et al., 2017; Cruz et al., 2019; Sarker and Oba, 2020). Our results suggest an alternative strategy for *S. parvula* where we observed an increase in both shoot and root xylem vessel area in plants exposed to long-term salt stress (given via 150 mM NaCl) (Figures 3 and 4) (Li et al., 2021). The salt-mediated expansion of xylem tissue in *S. parvula* was the second-most highly induced trait that responded by a higher fold change between control and salt-treated plants (Figure 8). This adaptive feature likely allows a higher bulk flow through the xylem to help *S. parvula* leaves maintain a cooler temperature while allowing uncompromised gas exchange compared to *A. thaliana* grown under saline conditions (Figure 5), despite the tradeoff of a higher risk of cavitation. Salt-induced increases to xylem diameter correlated to maintaining growth under salt stress has been reported for a few extremophytes (e.g. *Nitraria retusa* and *Atriplex halimus*) adapted to extreme salinities above seawater strength, but is not a common trait associated with halophytes (Boughalleb et al., 2009; Parida et al., 2016). The expansion of xylem area without the expansion of root or stem area that we observed as a salt responsive trait in *S. parvula* (Figure 3) is an atypical adaptation among plants resilient to salt stress (Olson and Rosell, 2013; Nassar et al., 2020). *S. parvula* seems to optimize effective use of water instead of simply reducing water loss at high salinities (Figures 5F and S9). Prioritizing effective water use has been proposed to be a better strategy for plant growth during water deficit stress than maximizing water conservation at the expense of photosynthetic capacity (Blum, 2009). This finding has implications for agriculture. If the availability of water can be ensured in agricultural systems even if the water source is brackish, the crops that are able to maintain relative water content (as observed for *S. parvula* in Figure 5), while allowing uncompromised transpiration and gas exchange, will be more resilient to environmental stresses.

### Adjustments to shoot architecture modulation follows root responses to cope with salt stress

*Schrenkiella parvula* root growth and development was less affected by increasing salt concentrations than in *A. thaliana*. *Schrenkiella parvula* primary root length increased while root fresh weight was only maintained under high salt conditions (Figures 2, S1 and S3), suggesting that salt causes a reallocation of resources toward deeper roots. This is also supported by the reduction in root hair length, total lateral root length and number (Figures 2 and S1). Given the natural environment in which the *S. parvula* ecotype Lake Tuz grows, the top layers of soil will dry first, causing the soil salt concentration to form a high to low gradient from topsoil to deeper layers in the rhizosphere (Tug et al., 2019). Thus, *S. parvula* can avoid relatively higher salinities by growing deeper to lower salt concentrations. The growth strategy exemplified by *S. parvula* is correlated with uncompromised growth dependent on modulating root to shoot vasculature and seemed to be efficiently coupled with leaf traits in *S. parvula*. Even though saline soils directly affect roots, its effects are also observed in shoots. If roots grow continuously in saline media, salts are often deposited in leaves. As plants age, salt-adapted plants either need to store salts in leaves and shed those whose capacity to hold excess salt is reached, or extrude excess salt via salt glands (Jennings, 1968; Dassanayake and Larkin, 2017). *Schrenkiella parvula* uses the first strategy by developing succulent leaves that are equipped with larger vacuoles which support salt sequestration. This trait appears to be facilitated by shifting the leaf cell population from a dominant diploid state to higher ploidy levels during prolonged salt stress (Figure 4). Indeed, the increase in leaf thickness, along with larger leaf area, was among the top traits with highest salt-induced fold-changes in *S. parvula* (Figure 8). Developing larger cells that can store excess salts within larger vacuoles enabled by endoreduplication has been reported for several other salt adapted extremophytes. It was proposed to be a key mechanism for surviving environmental stress (De Rocher et al., 1990; Barkla et al., 2018), while leaf succulence is one of the most common traits observed in halophytes (Jennings, 1968; Flowers and Colmer, 2008).

To maximize storage capacity via increasing the leaf surface area to volume, leaves are generally more terete in many extremophytes that have fully developed succulent leaves. This trait is often accompanied by the loss of leaf abaxial identity whereby the leaves become adaxialized (Ogburn and Edwards, 2013). Multiple genetic networks regulated by adaxially expressed HD-ZIPIII transcription factors have been shown to control leaf polarity (Du et al., 2018). *Schrenkiella parvula* leaves tend to acquire a more terete state with prolonged exposure to salt (Figure S7B). This phenotypic change is coincident with *S. parvula* leaves exhibiting reduced differentiation between adaxial and abaxial surfaces as leaf thickness increases with higher salt concentrations and a clear palisade layer at the adaxial surface is not present. However, this structural alteration does not affect the relative water content nor cause leaf curling or other visibly malformed vascular structures, as seen in adaxialized leaves of *A. thaliana* mutants (Figures 4, 5, and S7) (Du et al., 2018).

*Schrenkiella parvula* leaves are amphistomatous (possess stomata on the upper and lower leaf surfaces), in contrast to the *A. thaliana* leaves, which have more stomata on the abaxial surface (hyposomatous) (Figure 5B). Amphistomatous leaves are relatively rare in angiosperms (Nadeau and Sack, 2002; Drake et al., 2019) but in such leaves, the distance between stomata and the mesophyll is reduced allowing greater water-use efficiency due to increased CO_2_ conductance in the mesophyll thereby facilitating a higher relative photosynthesis rate (de Boer et al., 2016). Therefore, amphistomatous leaves are found in highly productive fast-growing herbs. However, when this trait is found in extremophytes such as *S. parvula* with trichomeless leaves arranged mostly with vertical leaf angles (Fig 1), it adds a tradeoff for water loss through transpiration as well as higher exposure to solar heat (Drake et al., 2019). Drake et al. (2019) hypothesized that to avoid desiccation in thick amphistomatous leaves, vein length per unit leaf area must be larger than in hypostomatous leaves of the same thickness. This anatomy would need to be accompanied by additional traits to facilitate an increase in transpirational flow via xylem development from root to shoot if leaf thickness increases as a response to salt stress. Additionally, having mostly vertical leaf angles with thick amphistomatous leaves is considered a key adaptive trait to reduce the effect of thermal radiation around midday while maximizing photosynthesis in the morning and late afternoon thereby lowering the risks of desiccation (King, 1997). Further, traits known to have evolved under shade avoidance such as internode elongation seems to be exapted in *S. parvula*, most likely to enhance transpirational cooling selected under environments that demand rapid growth amidst extreme environmental constraints (Pierik and Testerink, 2014).

Moderate to high salt is a persistent edaphic factor in the natural habitat of *S. parvula* (Helvaci et al., 2004; Tug et al., 2019). Such an environment would have selected for multiple traits that act synergistically to create an efficient lifestyle to survive stress. However, unlike many slow-growing extremophytes, *S. parvula* shows fast growth achieved with short life cycles, while enduring multiple environmental stresses. Often environmental stress resilience comes at the cost of yield reduction in crops and the drive for increasing yields has been prioritized over the need to develop resilient crops (Pardo and VanBuren, 2021). Increasing threats to global agriculture due to climate change necessitate a change in our priorities for crop development (IPCC, 2021). *Schrenkiella parvula* provides an excellent genetic model system that illustrates growth optimization over growth inhibition to cope with environmental stresses that we can explore to find innovative genetic architectures suitable and transferable to develop resilient crops.

## Materials and Methods

### Plant growth conditions

*Schrenkiella parvula* (Lake Tuz ecotype) and *A. thaliana* (Col-0 ecotype) were grown on soil, plates or in a hydroponic system as described in Wang et al., (2019), Pantha et al.,(2021), and Conn et al., (2013). Soil grown plants were used for reproductive trait assessments. Plate grown plants were used for root growth assessments and thermal tolerance assays. Hydroponically grown plants were used for shoot physiological assessments and anatomical characterization. See supplementary method S1 for details.

### Root growth assays

Five-day-old seedlings germinated on plates were transferred to 1/4x MS supplemented with 100 and 150 mM NaCl and kept at a photoperiod of 12 hr light/12 hr dark for 13 days. Additionally, 4-day-old seedlings of *A. thaliana* and *S. parvula* grown at a photoperiod of 16 hr light/8hr dark and on 1/2x MS agar plates were transferred to 1/2x MS plates supplemented with 125, 175, and 225 mM NaCl and monitored for 6 days. These plates were imaged for further processing using ImageJ (Ferreira and Rasband, 2012) to quantify primary root length, lateral root number and density, total and average lateral root length, and root hair length (Figures 2 and S2). See supplementary method S1 for details.

### Halotropism assay

Seedlings were grown on 1/2x MS medium solidified with 1.2% (w/v) Phyto agar plates held at an 85° incline as described in Galvan-Ampudia et al., (2013). After 5 days, the agar medium 1 cm below the root tips, was removed and replaced by 1/2x MS agar supplemented with 200 mM NaCl and seedlings were allowed to grow for another 5 days (Figure S4).

### Germination assay

Sterilized stratified seeds were germinated on 1/4x MS (Figures 7A, D, E and F) or 1/2x MS (Figures 7B and C) supplemented with different concentrations of NaCl, KCl, LiCl, or H_3_BO_3_. Final percent germination was recorded 3 days and germination curves recorded for up to 12 days after stratification. Seeds were counted as germinated if radicle emergence was observed. To check the viability of ungerminated seeds, *S. parvula* seeds that failed to germinate on high NaCl or KCl plates were transferred back to 1/4x MS or 1/2x MS and monitored for up to two weeks.

### Thermal stress tolerance assays

Five-day-old seedlings were subjected to heat stress at 38 °C for 6, 18 and 24 hr; chilling stress at 4 °C for 24 hr; or freezing stress at 0 °C for 12 hr followed by a recovery time of 5 days at 23 °C. Seedlings were considered to have survived if roots continued to grow without visible chlorosis of the cotyledons after the recovery period monitored for another 5 days.

### Quantification of leaf traits

Stomatal conductance and CO_2_ assimilation rate were measured using the sixth or seventh leaf from the shoot tip of hydroponically grown plants in LiCor 6400-40 leaf chamber attached to a 6400XT gas analyzer (LiCor Inc, Lincoln Nebraska). Measurements were taken after acclimation to 400 μmol photons m^-2^ s^-1^, 400 μL L^-1^ CO_2_ at 23 °C and an air flow rate of 200 µmol s^-1^ until a steady-state photosynthesis rate was attained. Each leaf was measured three times and three plants were used for each treatment.

Four-week-old hydroponically grown plants subjected to 150 mM NaCl or control treatments were used for the following leaf measurements. To quantify total leaf area, all leaves from selected plants were scanned (Perfection V600 scanner, Epson, Suwa, Nagano, Japan) and analyzed with ImageJ (Ferreira and Rasband, 2012). To obtain the relative water content (RWC), newly harvested leaves were weighed before and after submerging in water for 24 hr (to get fresh and turgid weights respectively) followed by drying until dry weights were obtained. RWC was calculated as (fresh weight - dry weight)/(turgid weight - dry weight). Leaf temperature was measured on intact leaves of 4-week-old plants between ZT3 to ZT4 (Zeitgeber Time) every day for 7 days using thermal imaging (E6, FLIR, Wilsonville, Oregon). Leaf temperatures were averaged across all leaves for each plant.

### High-throughput phenotyping of morphometric and physiological traits

*A. thaliana* and *S. parvula* soil grown plants were phenotyped for compactness, leaf water status, leaf area, and chlorophyll fluorescence using a PSI PlantScreen™ Compact system (Qubit Phenomics, PSI). See supplementary method S1 for details.

### Anatomical and flow cytometry analyses

Sections obtained from young roots (1 cm from the tip), mature roots (2 to 5 cm from the root and shoot junction), stems (fourth, fifth, and sixth internodes from the shoot tip) and leaves (fourth, fifth, and sixth from the shoot tip) from 8-week-old plants were used for anatomical trait characterization. The fifth and sixth leaves from the shoot tip were used for flow cytometry analysis as described in Galbraith et al., (1983). We counted the number of stomata stained with propidium iodine on both hydroponically- and soil-grown plants and on adaxial and abaxial surfaces at the same developmental stage and treatment duration. See supplemental method S1 for details.

### Reproductive trait quantification

Three-week-old soil grown plants were treated with an incremental increase of 50 mM NaCl every 2 days until the final concentration was reached to 150 mM NaCl treatments and maintained at 150 mM NaCl for the remainder of the experiment under either a long-day (16 hr light/ 8 hr dark) or short-day (12 hr light/12 hr dark) photoperiod. Flowering time was recorded when the first open flower was observed. Total numbers of flowering events, including all developing, mature, and aborted flowers, as well as fully developed siliques were counted separately for each plant throughout its lifetime. Fully developed siliques were counted and harvested for imaging from 5-week-old plants that were given 150 mM NaCl or control treatments for an additional 2 weeks.

### Computational analysis

The data used to compare the basal expression of wax biosynthesizing genes between *A. thaliana* and *S. parvula* were retrieved from NCBI-SRA database, accession# SRX877979 and SRX877980, respectively (Oh et al., 2014). The expression profiles were visualized using Integrated Genomics Viewer (v2.6) (Thorvaldsdóttir et al., 2013). The copy numbers and transposition states of PhyB/D orthologs were determined using CLfinder-OrthNet pipeline as described by Oh and Dassanayake, (2019). For principal component analysis (PCA) given in Fig 8, as traits were measured in different units and scales, the measurements for each trait were first normalized by either subtracting the mean (for anatomical traits) or dividing by the mean (for physiological traits). The normalized data were subsequently analyzed with prcomp in R.

## Acknowledgement

This work was supported by the US National Science Foundation/US-Israel Binational Science Foundation award NSF-BSF-IOS-EDGE 1923589/2019610, NSF-MCB-1616827, US Department of Energy BER-DE-SC0020358, Next-Generation BioGreen21 Program of Republic of Korea (PJ01317301), and the Goldinger Trust Jewish Fund for the Future awards. Graduate students K.T., G.W., P.P., and C.W. were supported by an Economic Development Assistantship and undergraduate students J.C.G. and M.G.M. were supported by the President’s Future Leaders in Research Program at Louisiana State University (LSU). H.L. was sponsored by the China Scholarship Council (CSC) through a Sino-Dutch Bilateral Exchange Scholarship. C.T. and Y.Z. were funded by the European Research Council (ERC) under the EU Horizon 2020 Research and Innovation programme (grant #724321). We acknowledge the productive discussions led by Drs. Hans Bohnert and John Cheeseman at the University of Illinois at Urbana-Champaign (UIUC) that prompted us to initiate this study. We thank the LSU High Performance Computing facility for providing computational resources; Dr. Ying Xiao in the Shared Instrumentation Facility at LSU for assistance with microscopy imaging; undergraduate students Stephanie Presedo at LSU and Rebekah Munaretto, Michael Pettineo, and Aditya Ravindra at UIUC for assisting with plant phenotyping; Jasper Lamers from WUR-PPH for providing the script for automated root edge detection; and Aliza Finkler and Guilia Meshulam at Tel Aviv University for providing technical help with the phenomics study.

## Author contributions

K.T., P.P., G.W., and N.K. conducted the main experiments and analyzed data as major contributors to the overall project; C.W., D-H.O., N.D., H.L., H.H., and P.A. conducted additional experiments and analyzed data; undergraduate students J.C.J., R.K., M.G.M., A.C., D.T., and high school student C.C. conducted complementary experiments supervised by K.T., P.P., G.W., D-H.O. and C.W.. M.D., C.T., and S.B. designed independent complementary experiments; supervised students in their labs; and contributed to data interpretation. D.L., Y.Z., and G.S. provided additional student supervision and assisted with data generation for selected experiments. K.T., P.P., G.W., S.B., and M.D. wrote the manuscript. K.T., P.P., G.W., N.K., D-H.O., C.W., P.A., M.F., J.C.L., A.S., D.L., P.F., C.T., S.B., and M.D. provided critical reviews. M.D. conceptualized and supervised the overall project.

---

**Figure S1.**
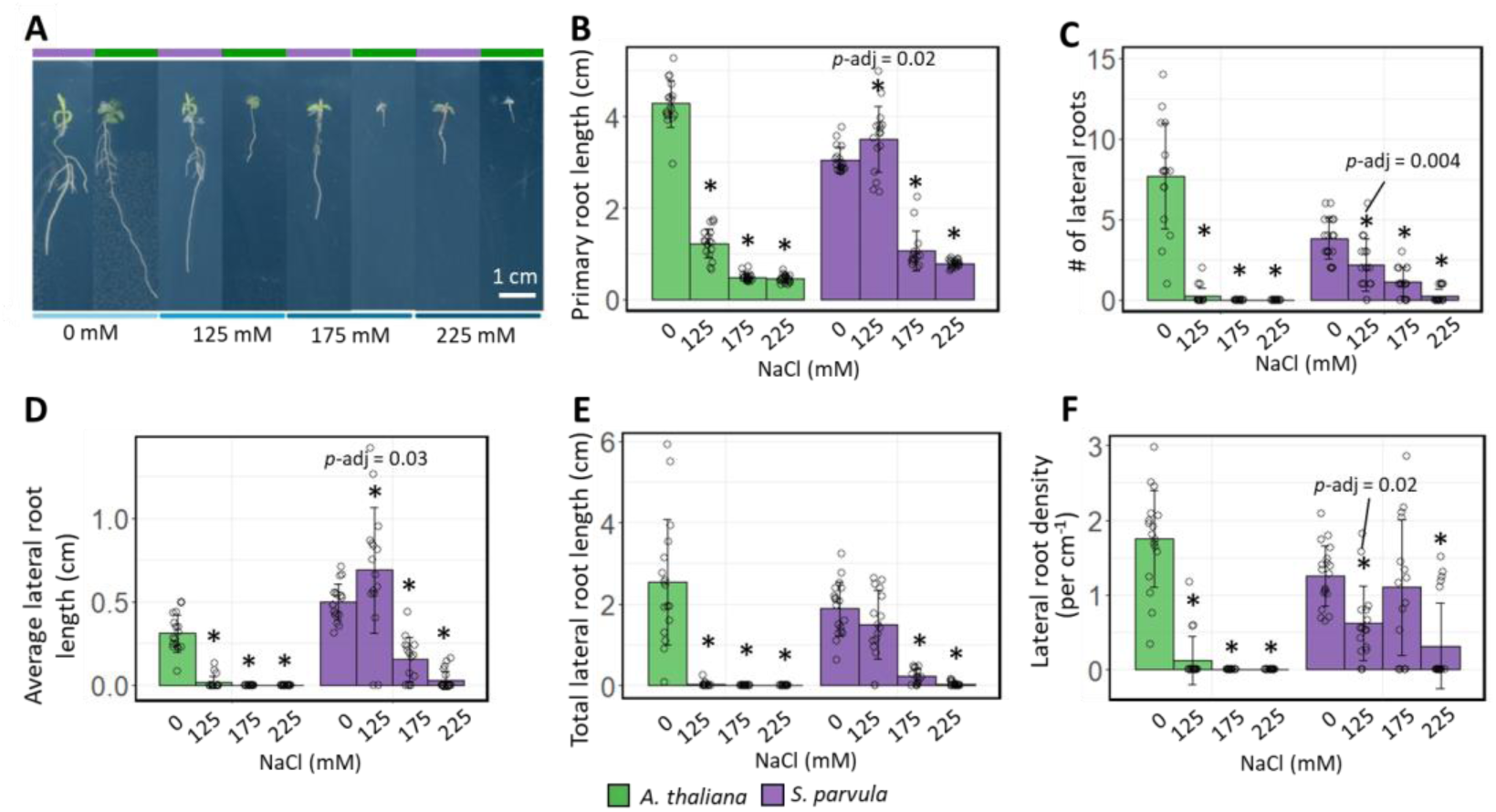
Effects of NaCl stress on root growth of *Schrenkiella parvula* and *Arabidopsis thaliana* under long days and higher nutrient conditions. [A] Root growth of 10-day-old seedlings under indicated concentrations of NaCl. Quantification of [B] primary root growth, [C] number of lateral roots, [D] average lateral root length, [E] total lateral root length, and [F] lateral root density. Asterisks indicate significant difference (p-adj ≤ 0.05) between the treated samples and their respective control samples, determined by two-way ANOVA. Data are represented as mean ± SD (n = 15 to 20). Open circles indicate the number of plants used for each experiment.

**Figure S2.**
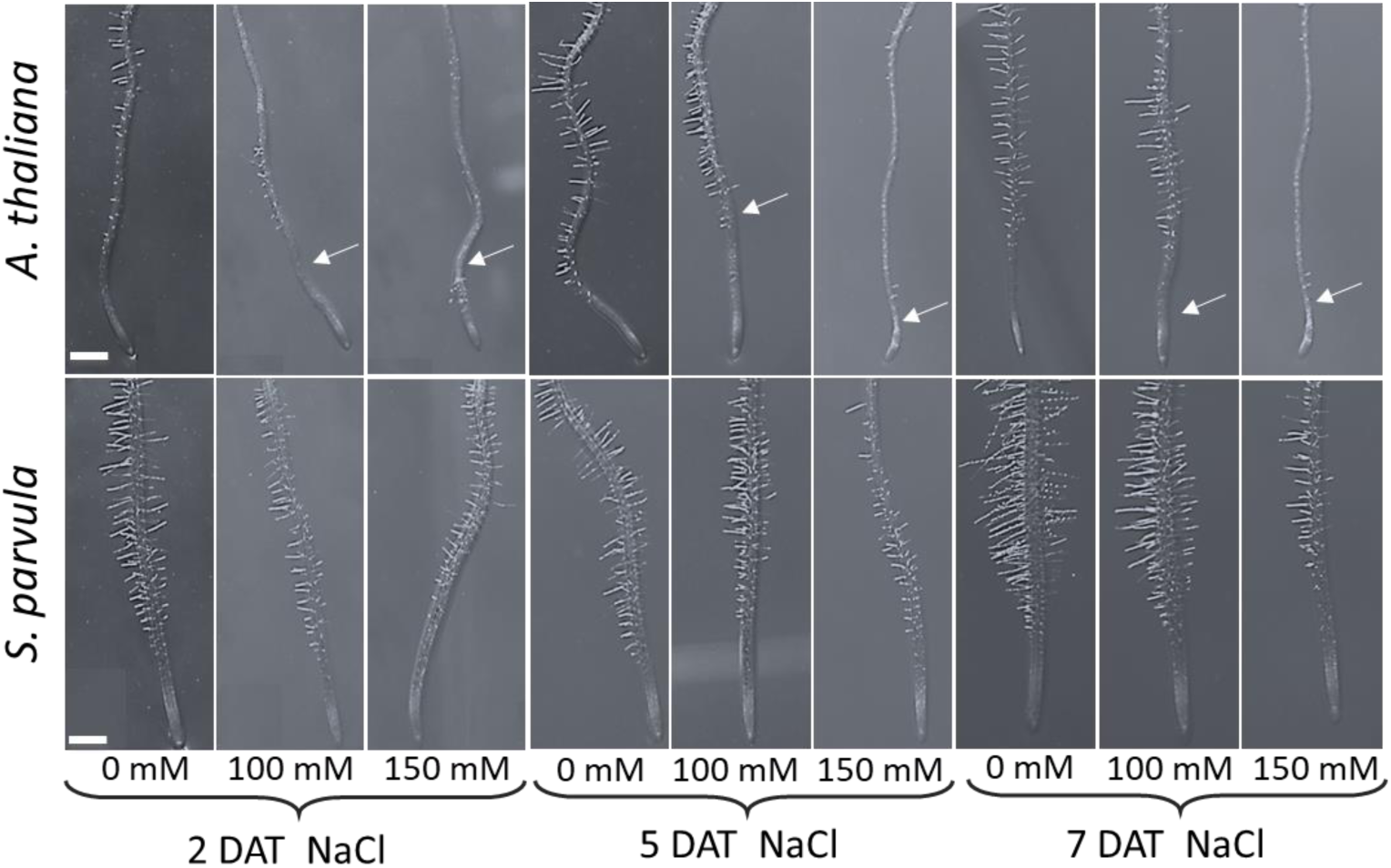
Effects of NaCl stress on root hair growth in *Schrenkiella parvula* and *Arabidopsis thaliana*. White arrows indicate root tip positions when the 5-day-old seedlings were transferred to the indicated salt concentrations. Note that the white arrows were absent in *A. thaliana* and *S. parvula* control because the position of the root tip at the time of transferring were out of frame when we imaged the root tip region. Scale bars are 0.5 mm. DAT, Days after treatment.

**Figure S3.**
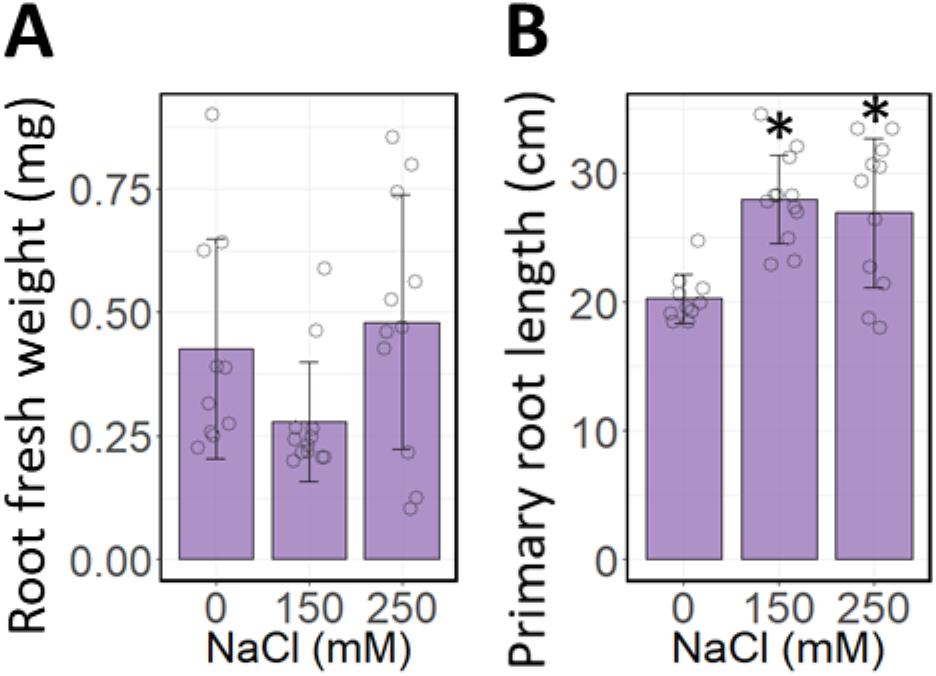
Long term effects of NaCl stress on the root growth of *Schrenkiella parvula*. [A] Root fresh weight and [B] primary root length of 12-week-old hydroponically grown *S. parvula* plants. Four-week-old plants were subjected to 150 mM NaCl for an additional 4 weeks and the treatment was continued or increased to 250 mM NaCl for another 4 weeks. The plants were grown at 100-150 μmol m^-2^ s^-1^ photosynthetic photon flux density with a 12 hr light/12 hr dark photoperiod. Asterisks indicate significant difference (*p* ≤ 0.05) between the treated samples and their respective control samples, determined by Student’s *t*-test. Data are represented as the mean ± SD (n = 9). For each plot, an open circle indicates the measurement from each plant.

**Figure S4.**
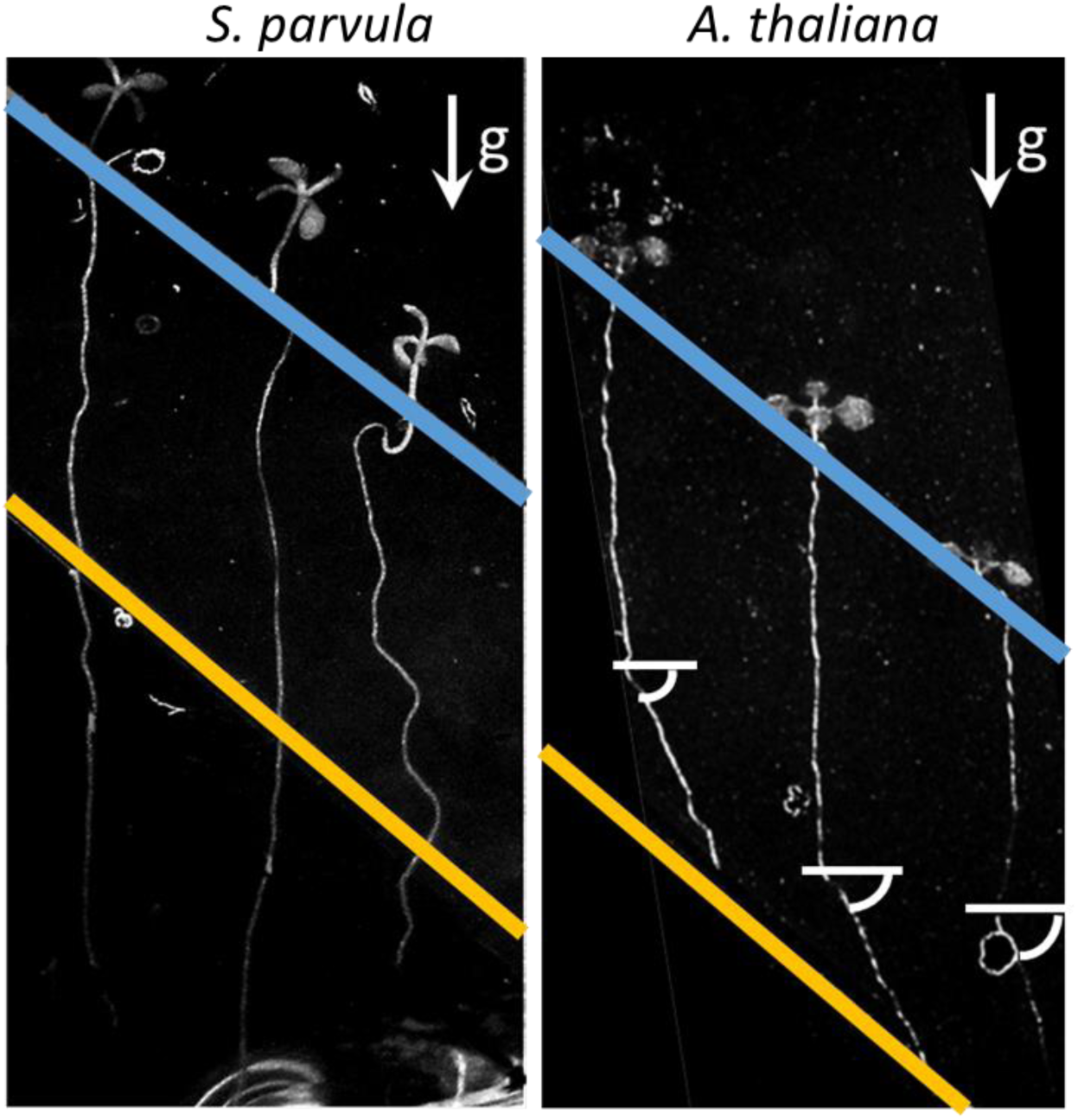
Halotropism assay at 200 mM NaCl for *Schrenkiella parvula* and *Arabidopsis thaliana*. The blue lines indicate the position of seeds sown. The yellow line marks the separation between 1/4x MS media with 0 mM (top) and 200 mM NaCl (bottom). The bent root angle is indicated by curved white lines. g, gravity axis.

**Figure S5.**
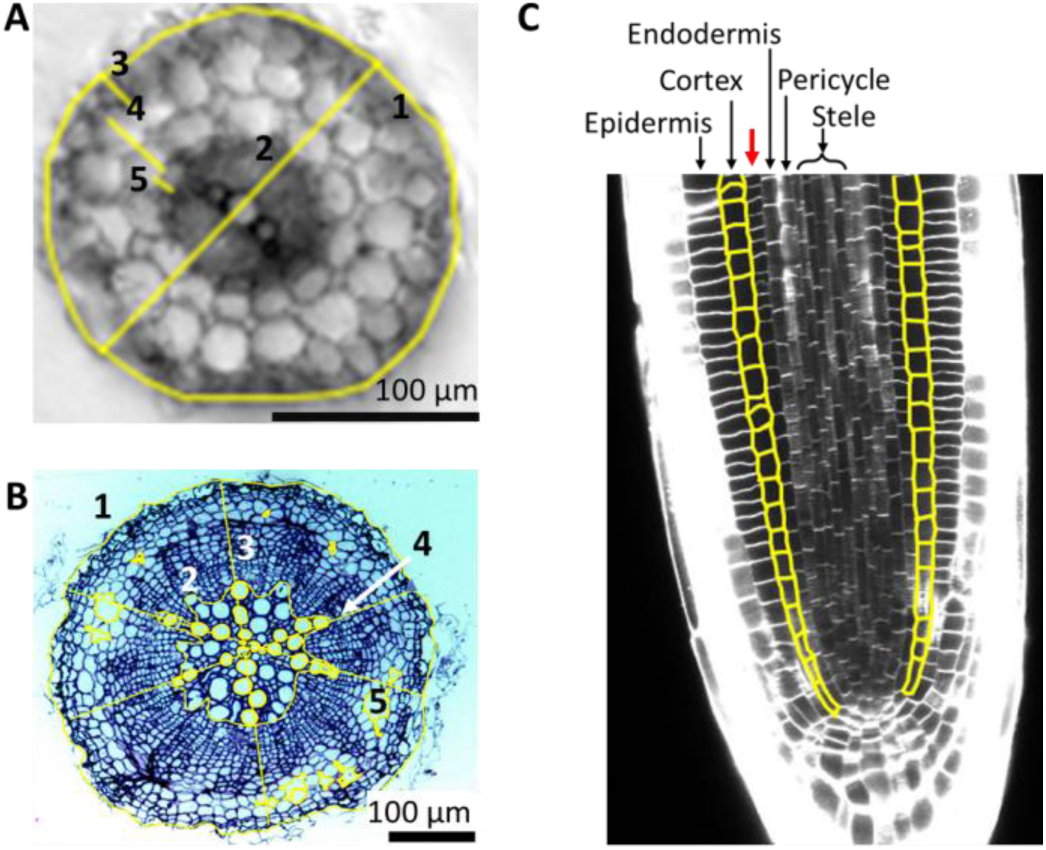
Root anatomy of *Schrenkiella parvula*. [A] Transverse section of the young root taken at 1-3 cm from the root tip. [1] Young root area; [2] root diameter; [3] epidermis thickness; [4] cortical layer thickness; [5] endodermis thickness. [B] Transverse section of the mature root taken between 2-5 cm below the root-shoot junction. [1] Mature root area; [2] xylem/root area; [3] Area per vessel; [4] # vessels/root diameter; [5] Air space/root area. [C] Longitudinal section of young root tip from 7-day-old seedlings. The cortex layer is highlighted in yellow. The red arrow indicates the additional cell layer.

**Figure S6.**
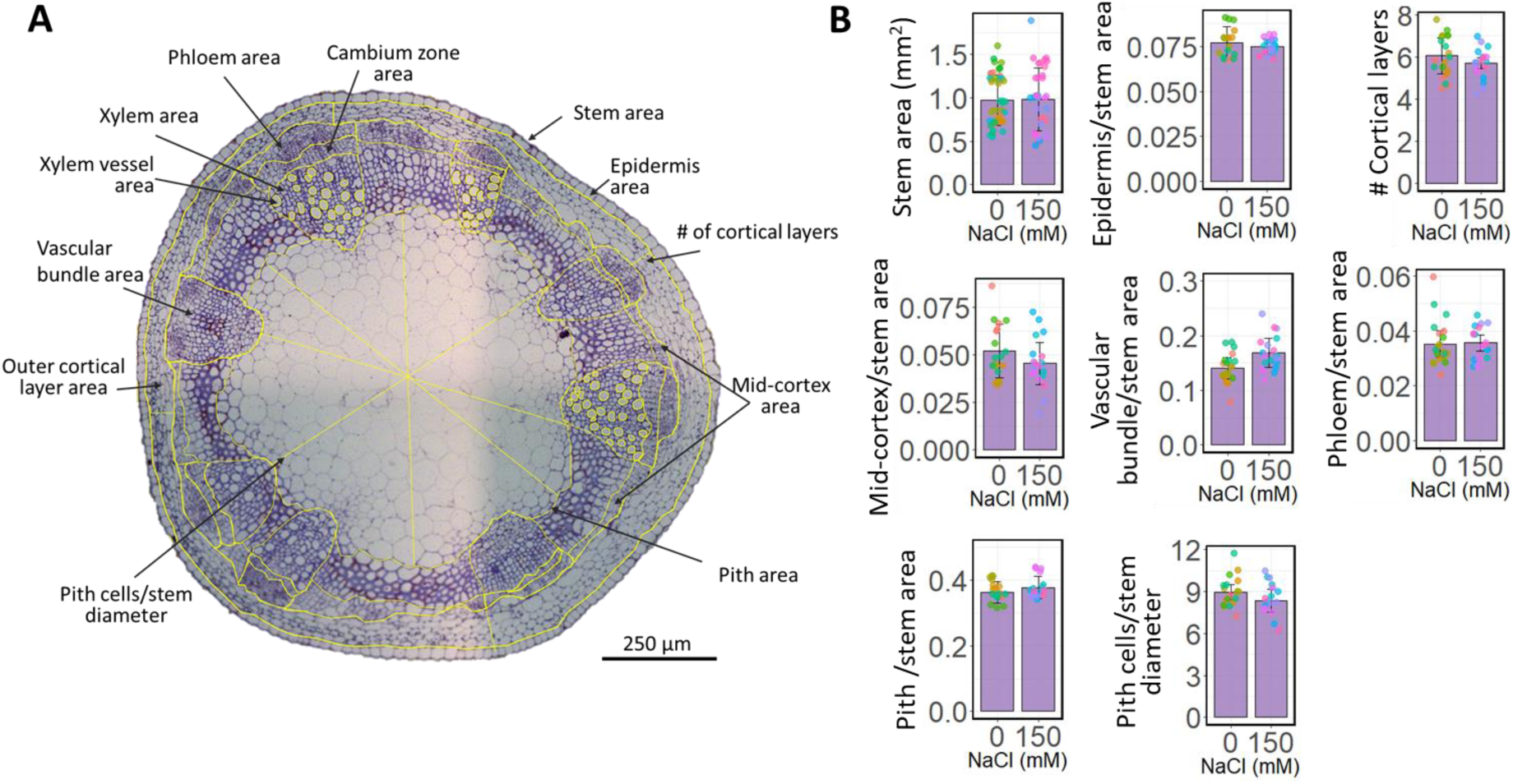
Stem anatomy of *Schrenkiella parvula*. [A] Transverse section of *S. parvula* stem (between 4th, 5th, and 6th internodes from the shoot meristem) with cell layers marked in yellow lines for trait quantification. [B] Measurements of stem anatomical features indicated in [A]. A minimum of 20 sections from 5-11 plants were used for each measurement. Asterisks indicate significant difference (*p* ≤ 0.05) between the treated samples and its respective control samples, determined by Student’s *t*-test. Data points represent individual cross-sections and colors represent individual plants.

**Figure S7.**
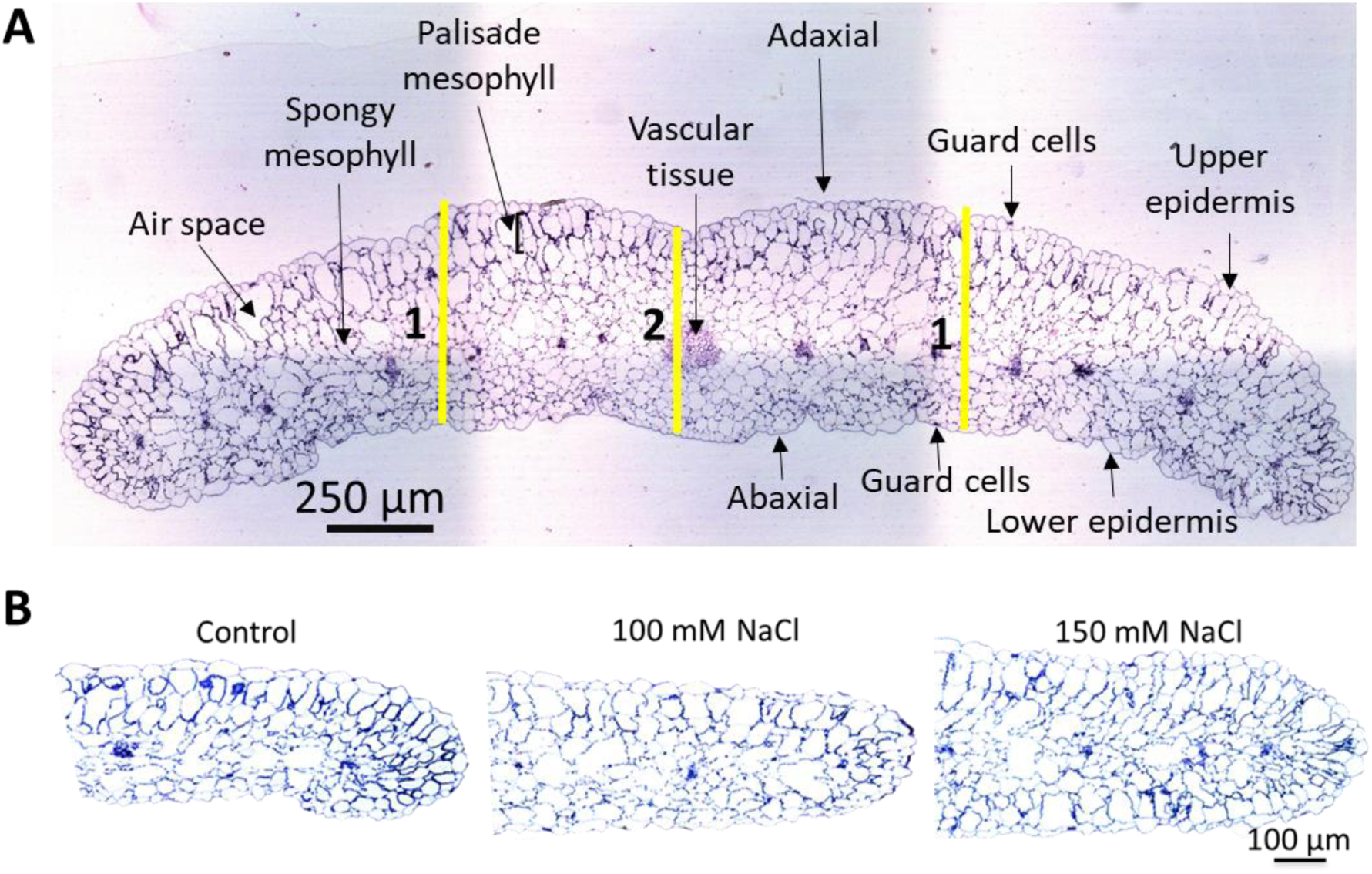
Leaf structures of *Schrenkiella parvula*. [A] Transverse section of *S. parvula* leaves. [1] leaf thickness and [2] midrib thickness. [B] Transverse sections of *S. parvula* leaves treated with indicated concentration of salts. All sections represent the fifth or sixth leaf from the root-shoot junction in 8-week-old plants that were treated for 4 weeks.

**Figure S8.**
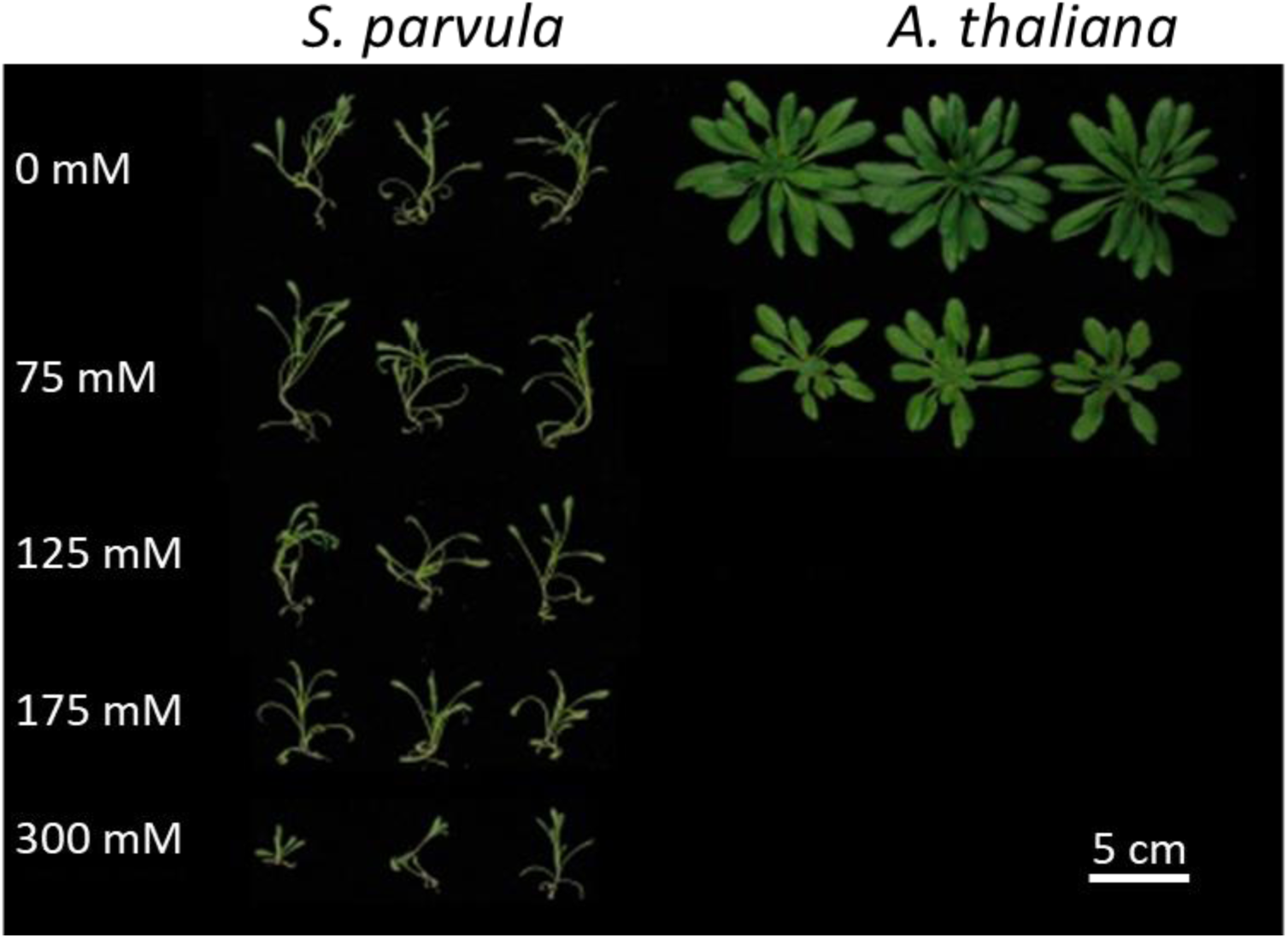
Effects of salt on growth of 4-week-old *Arabidopsis thaliana* and *Schrenkiella parvula* treated for an additional 2 weeks at 22 °C under a 12 hr light/12 hr dark photoperiod in soil.

**Figure S9.**
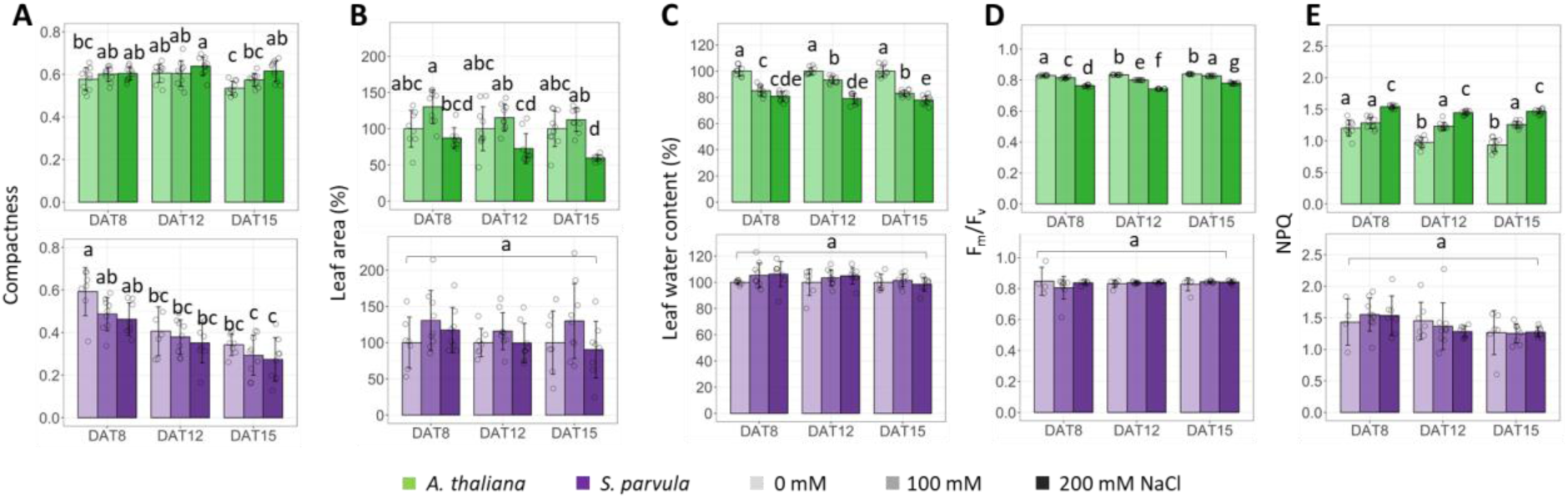
Effects of NaCl stress on *Arabidopsis thaliana* and *Schrenkiella parvula* morphometric and physiological traits. Soil-grown, 8-day-old *A. thaliana* and 12-day-old *S. parvula* were exposed to the indicated salt concentrations at 21 °C, 16 h light/8 h dark photoperiod. The Measurements were taken on day 8, 12, and 15 after the start of the salt treatments. [A] Compactness (the ratio between the rosette (or leaf) area and the rosette (or leaf) convex hull area); [B] Leaf area; [C] Leaf water content; [D] Maximum quantum efficiency of PSII; and [E] Non-photochemical quenching were measured/estimated using the Qubit/PSI PlantScreen™ Compact System. For leaf area and leaf water content, values are percent of control (0 mM NaCl) for each time point. Letters denote significant changes at *p*-adj ≤ 0.05 (one-way ANOVA with post-hoc Tukey’s test). Data are mean ± SD (n = 10). Open circles indicate the individual measurements obtained from each plant. DAT, days after treatment.

**Figure S10.**
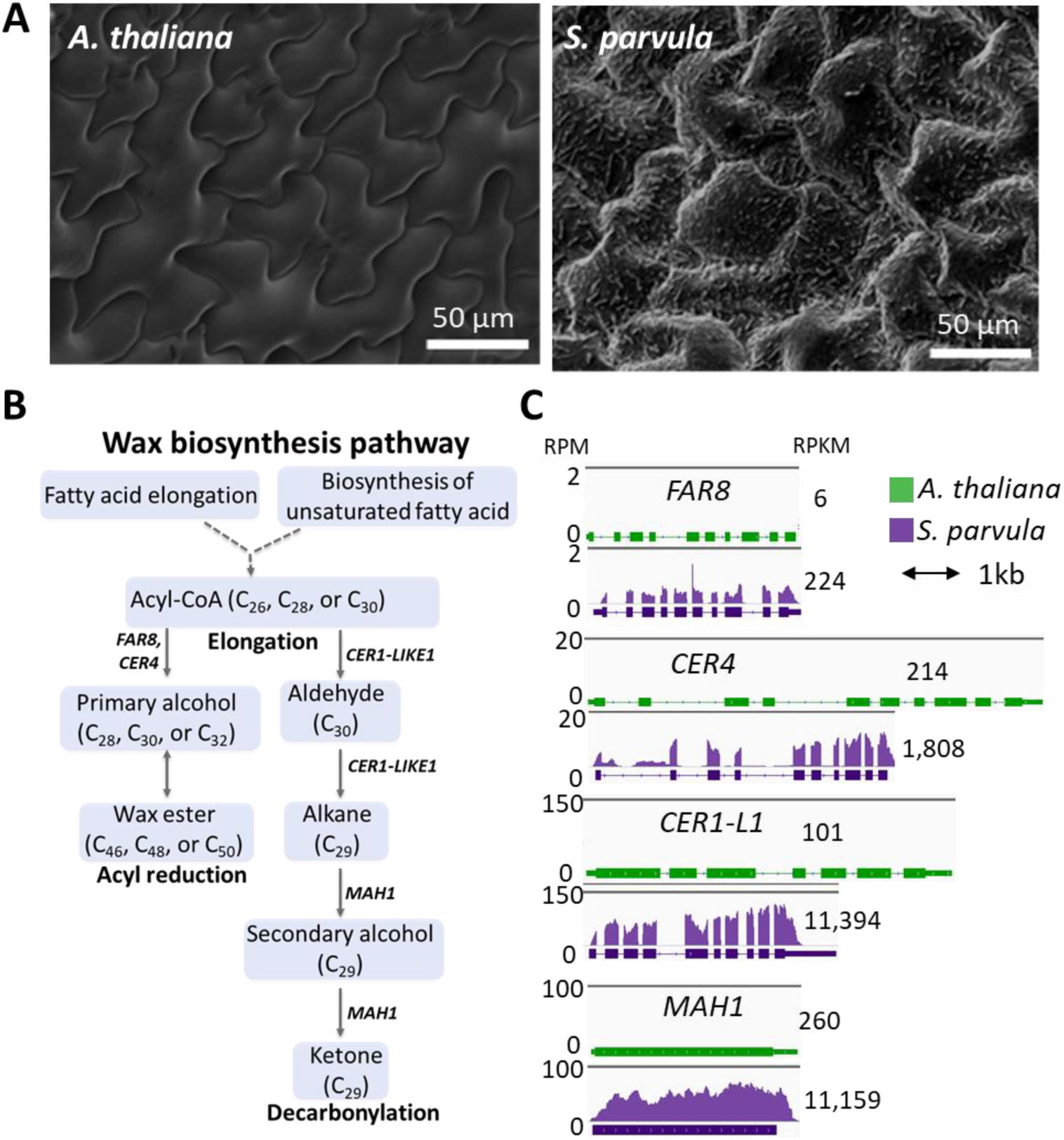
Leaf surface and the basal expression of genes involved in wax biosynthesis in *Arabidopsis thaliana* and *Schrenkiella parvula*. [A] Scanning electron micrographs contrasting *A. thaliana* and *S. parvula* leaf surfaces. [B] Major wax biosynthesis pathway and [C] wax biosynthesis genes that exhibited significantly different basal expression between *A. thaliana* and *S. parvula*.

**Figure S11.**
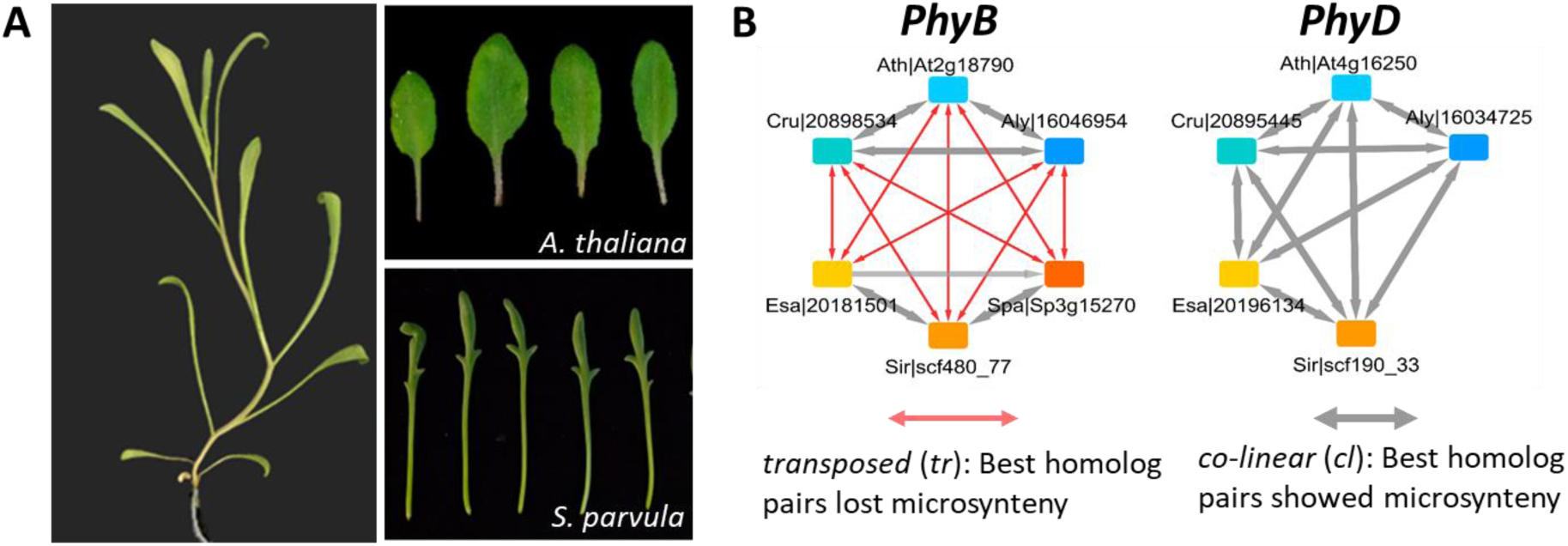
The potential influence of phytochrome family of genes on *Schrenkiella parvula* leaf and growth form. [A] Elongated internode and leaf petiole in *S. parvula* compared to *A. thaliana*. [B] *PHYD* is absent in the *S. parvula* genome as illustrated using OrthNet representing evolutionary histories of orthologous gene groups derived from six Brassicaceae genomes: *Arabidopsis lyrata* (Aly, version 1.0), *Arabidopsis thaliana* (Ath, v. ‘TAIR10’), *Capsella rubella* (Cru, v. 1.0), and *Eutrema salsugineum* (Esa, v. 1.0), *Sisymbrium irio* (Sir, v. 0.2), and *Schrenkiella parvula* (v. 2.0). Nodes are color-coded according to the species. Edges show either co-linear (cl) or transposed (tr) properties.

**Figure S12.**
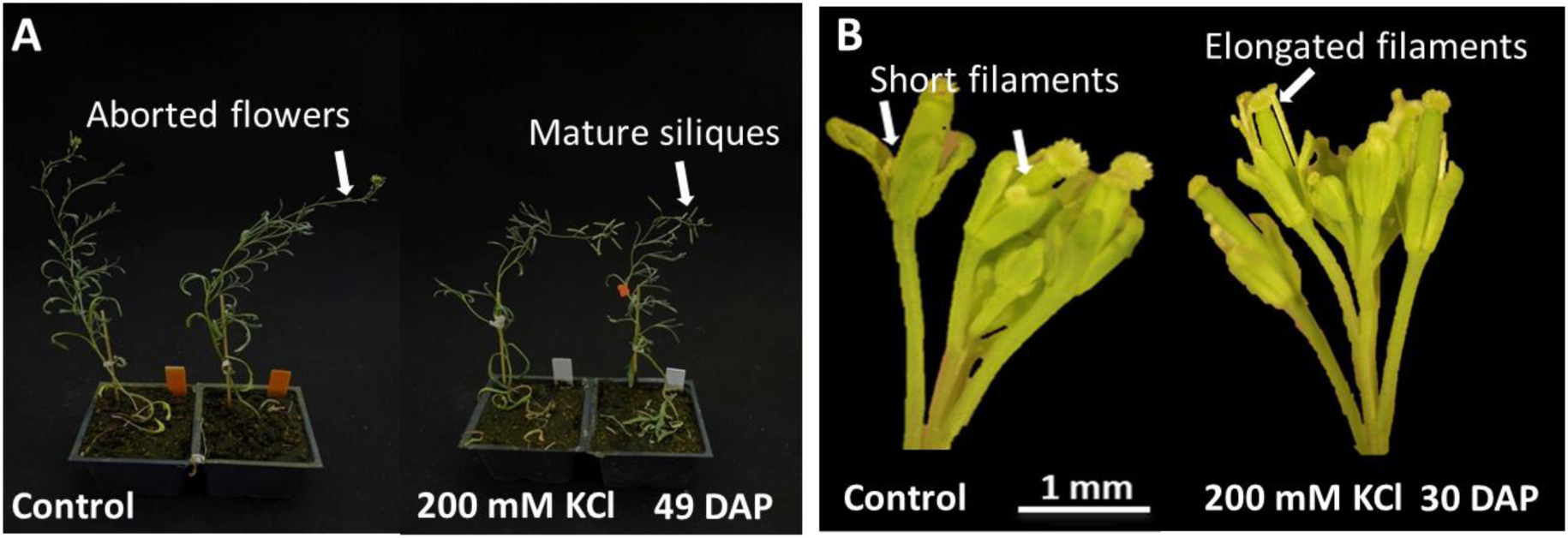
Effects of KCl on reproductive traits of *Schrenkiella parvula*. [A] Plants grown under control conditions generally flower late and the first few flowers are subsequently aborted. KCl-treated plants flower earlier and the first flowers develop into mature siliques. [B] Early flowers in KCl treated plants develop longer filaments compared to control plants. DAP, days after planting.

**Figure S13.**
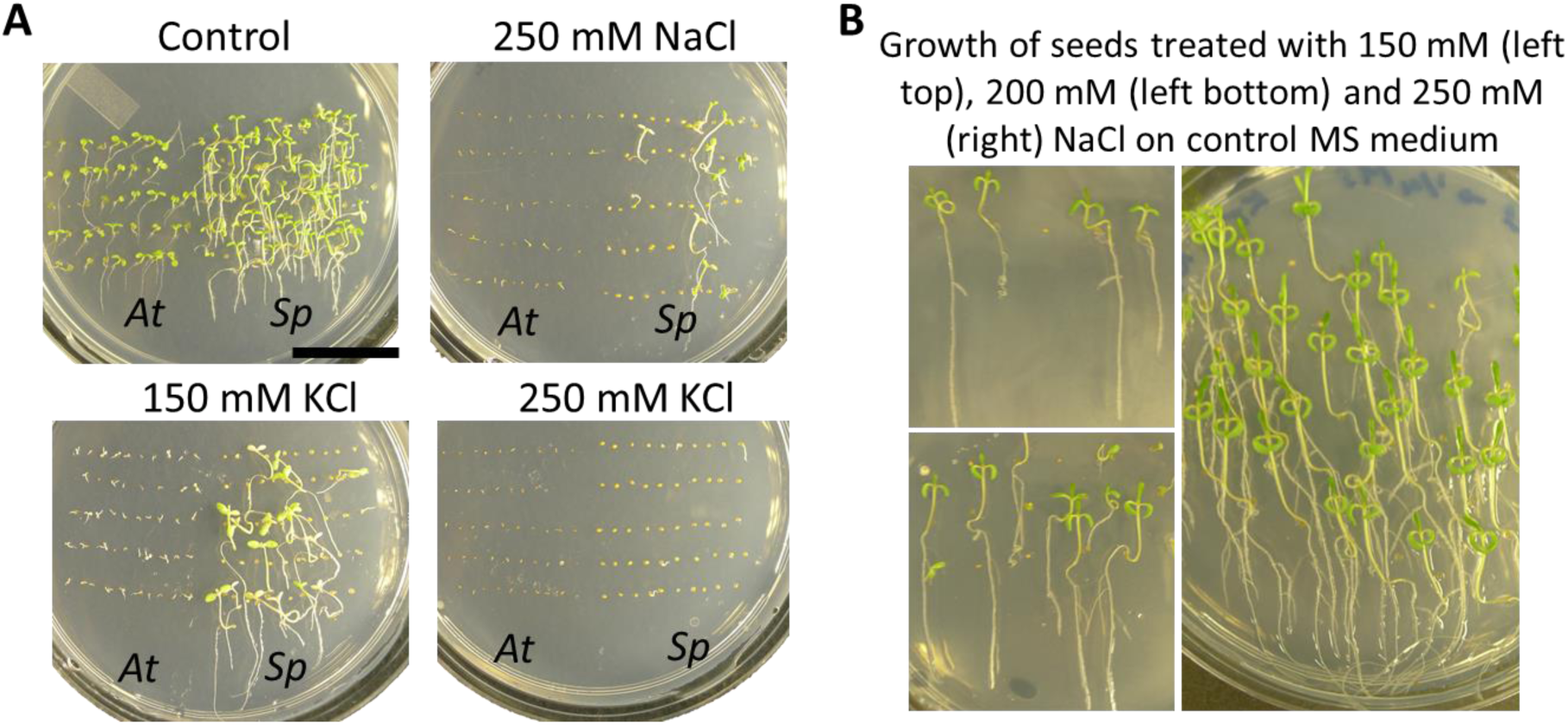
Effect of salt on *Arabidopsis thaliana* and *Schrenkiella parvula* inducing barriers for seed germination and seedling establishment. [A] *S. parvula* (*Sp*) and *A. thaliana* (*At*) seed germination and growth on NaCl- and KCl-supplemented MS medium. Scale bars are 2 cm. [B] Growth of ungerminated *S. parvula* seeds treated with different concentrations of NaCl (as shown in Figure 7) after transferring to control MS medium.

**Figure S14.**
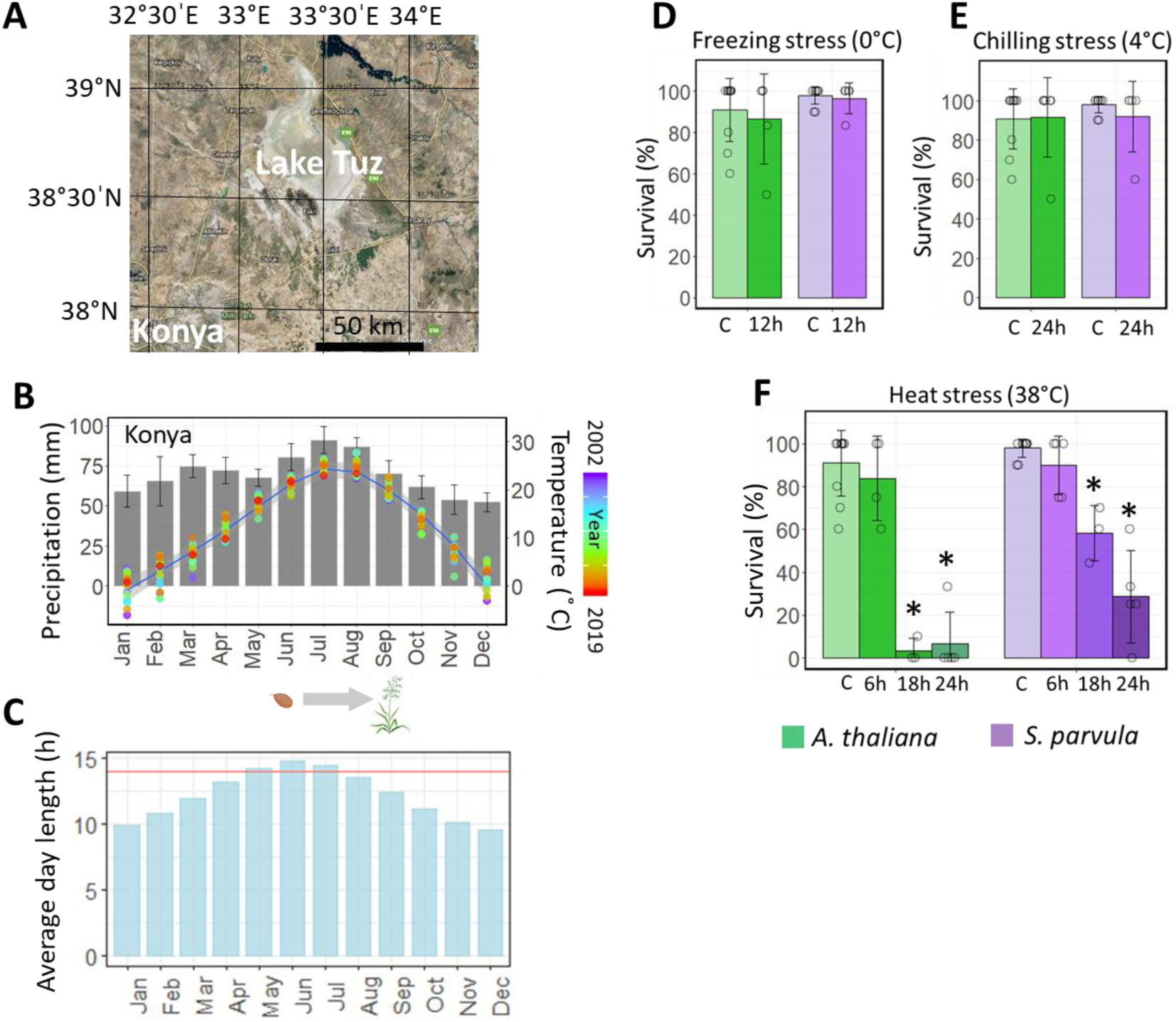
Survival of *Schrenkiella parvula* in the climate conditions found in Lake Tuz region. [A] Lake Tuz location from Google Earth; [B] Precipitation and temperature recorded in Konya, Turkey (NOAA, 2019). *S. parvula* growing season is from April/May to August/September (Tug et al., 2019). Bars indicate precipitation and lines represent temperature. [C] Average day length recorded in Konya per month. Red line indicates the day length used for typical laboratory growth assays for *S. parvula*. [D-F] Survival rate of *A. thaliana* and *S. parvula* seedlings under [D] freezing stress, [E] chilling stress, and [F] heat stress. Five-day-old seedlings were subjected to temperature treatments and data were recorded 5 days after recovery. Asterisks indicate significant difference (*p* ≤ 0.05) between the treated samples and their respective control samples (student’s *t*-test). Data are mean ± SD (n = 4). Each replicate plate contained 3-6 plants. Open circles indicate the biological replicates.

