## Supplemental method for "Balancing growth amidst salinity stress – lifestyle perspectives from the extremophyte model *Schrenkiella parvula*"

**Supplemental methods:**

**Plant growth conditions**

**Plate-grown plants.** *Schrenkiella parvula* (Lake Tuz ecotype) and *A. thaliana* (Col-0 ecotype) seeds were surface-sterilized, stratified for 5-7 days at 4 °C, and germinated on Murashige and Skoog (MS) medium (Murashige and Skoog, 1962) as described by Pantha *et al.* (2021). We used 1/4x (Figures 2, 7A, D, E, F, and S13) or 1/2x (Figures 7B, C and S1) MS plates. The plates were incubated at 23 °C with a light intensity of 100 to 150 µmol $m^{-2}s^{-1}$ photosynthetic photon flux density under a 12 hr/12 hr (Figure 2), or 14 hr/10 hr (Figure 7A, D, E, and F) or 16 hr/8 hr (Figures 7B and C; S1) photoperiod. Plate grown plants were used for root growth assessments and thermal tolerance assays.

**Hydroponically-grown plants.** Plants were grown hydroponically in 1/5x Hoagland's solution as described by Conn et al. (2013). Four-week-old plants were subjected to NaCl treatments for an additional 4 weeks unless otherwise indicated in the figure caption. These plants were used for shoot physiological assessments and anatomical characterization.

**Soil grown plants.** Plants were grown on soil as described by Wang et al., (2019) with a 14 hr light/10 hr dark photoperiod, 100 to 130 µmol $m^{-2}$ $s^{-1}$ light intensity, at 22 to 24°C unless otherwise indicated. These plants were used for reproductive trait assessments.

**Root growth assays**

Five-day-old seedlings germinated on plates were transferred to 1/4x MS supplemented with 100 and 150 mM NaCl and kept at a photoperiod of 12 hr light/12 hr dark for 13 days. These plates were imaged for further processing using ImageJ (Ferreira and Rasband, 2012) to quantify primary root length, number of lateral roots, and lateral root density (Figure 2). Root hair was imaged using a microscope (Zeiss Neolumar S 1.5 X FWD 30 mm) and the average length of the 10 longest root hairs were used to quantify root hair length for each plant (Figures 2D and S2). Additionally, 4-day-old seedlings of *A. thaliana* and *S. parvula* grown at a photoperiod of 16 hr light/8hr dark and on 1/2x MS agar plates were transferred to 1/2x MS plates supplemented with 125, 175, and 225 mM NaCl and monitored for 6 days. The primary root length, average and total lateral root length, lateral root number and density from 10-day-old seedlings were quantified. The plates were scanned at 400 dpi (Epson perfection V800 scanner, Suwa, Nagano, Japan), and pre-traced by an automated script based on root edge detection (https://github.com/jasperlamers/RootFinder/tree/main). Manual correction was performed with an ImageJ plugin, Smartroot (Lobet et al., 2011). Total lateral root length was the sum of length from all lateral roots in each plant. Average lateral root length was calculated by taking the total lateral root length divided by lateral root number. Lateral root density was obtained from dividing the number of lateral roots by the primary root length.

**Flow cytometry analysis**

The fifth and sixth leaves from the shoot tip of hydroponically grown plants were used for the extraction of protoplasts as described in Galbraith et al.,(1983). A flow cytometer (FACScan, BD Biosciences, San Jose, CA) with a 15 mW 488 nm argon-ion laser configured for propidium iodide fluorescence measurements were used for cell ploidy detection. A total of 15,000 cells per sample from three biological replicates were acquired and analyzed in the form of DNA ploidy histograms (Cellquest Pro software, BD Biosciences, San Jose, CA on a Macintosh G5 workstation, Apple Computer, Cupertino, CA).

**High-throughput phenotyping of morphometric and physiological traits**

*A. thaliana* and *S. parvula* seeds were surface-sterilized with 50% bleach for 5 min then rinsed 4 times with distilled water. Seeds were sown onto MS agar plates (4.4 g $L^{-1}$ MS salts, 0.5 g $L^{-1}$ MES (pH 5.7), 2% (w/v) sucrose, 0.8% (w/v) agar (Duchefa Biochemie) and stratified at 4 °C in the dark for 7 d (*A. thaliana*) or 14 d (*S. parvula*) before transfer to the growth room (16 h light, 8 h dark, 21°C). After 4 d (*A. thaliana*) or 6 d (*S. parvula*), plants were transferred to soil (Even-Ari Green 7611, 70% peat, 30% perlite 1.5 kg m-3 osmocote 14-14-14), with 1 plant per PlantScreen™ Compact system specific pot (200 ml, 5.4 x 5.4 cm at the top). Salt treatment (0, 100, or 200 mM NaCl) was initiated at 4 d (*A. thaliana*) or 6 d (*S. parvula*) after transfer to soil. Plants were irrigated twice a week for the duration of the experiment.

Plants were moved to the PSI PlantScreen™ Compact system (Qubit Phenomics, PSI) at Tel-Aviv University, 3 d prior to the start of the experiment, at 9 (*A. thaliana*) and 11 (*S. parvula*) days after transfer to soil respectively. Leaf area and compactness were measured using Red Green Blue (RGB) images taken from above using a 5 MPX camera under 180 µmol photons $m^{-2}$ $s^{-1}$

light. Images were automatically corrected for Barrel distortion and were used for morphometric measurements based on computer-generated masks of plant outline from which the system calculated leaf area and plant convex hull area in $mm^2$. Compactness was calculated by the PSI PlantScreen™ Compact system software as the ratio between the rosette (or leaf) area and the rosette (or leaf) convex hull area. Leaf water status was estimated based on leaf temperature measured via a top-mounted long-wave infra-red high-resolution camera and imaging at 940 nm and 1450 nm, which was converted to leaf water status by the PSI PlantScreen™ Compact system software. Chlorophyll fluorescence was measured at room temperature based on the adaxial side of rosette leaves. The following protocol was used: Plants were dark-adapted for 15 min prior to chlorophyll fluorescence measurements. Minimal fluorescence ($F_0$) was measured using 10 µs 620 nm (red-orange) flashes, followed by an 800 ms, 2000 µmol photons $m^{-2}$ $s^{-1}$ (actinic light 2: cool white) saturation pulse for measurement of maximum fluorescence ($F_m$). Plants were then subjected to 6 min of 180 µmol photons $m^{-2}$ $s^{-1}$, with saturation pulses every 30 s. Peak fluorescence (FP) was measured prior to the first saturation pulse, $F_m\_L_n$ was measured based on the maximum fluorescence signal during each saturation pulse, and the steady-state fluorescence signal ($F_t\_L_{ss}$) and maximum signal ($F_m\_L_{ss}$) were measured before and during the last saturation flash. Variable fluorescence in the dark-adapted state ($F_v$) was calculated by $F_m$-$F_0$. Maximum quantum yield of PSII was calculated as $F_v/F_m$; NPQ was calculated as $(F_m-F_m\_L_n)/F_m$ (Photon Systems Instruments, 2019). All calculations were performed by FluorCam7 software 2.1 (Photon Systems Instruments).

**Root and shoot anatomical analysis**

**Fresh sections.** Freshly-cut root tips were stained with 0.05% (w/v) Toluidine blue, washed with deionized water, and mounted with glycerol for root tip imaging. Fresh cross sections from young roots (1 cm from the tip), mature roots (2 to 5 cm from the root and shoot junction), stems (fourth, fifth, and sixth internodes from the shoot tip) and leaves (fourth, fifth, and sixth from the shoot tip) from 8-week-old plants were used for anatomical trait characterization. Root and shoot cross-sections were stained with 0.05% (w/v) Toluidine blue and 0.01% (w/v) Safranin-O, respectively, for 30 secs to 1 min. All fresh samples were examined for anatomical traits under bright field illumination using a DM6B Upright Microscope (Leica,

Wetzlar, Germany, which features a Hamamatsu sCMOS camera, Japan).

**Fixed sections.** Plant cross sections of mature roots, stems, and leaves were prepared by fixing tissues overnight at 4 °C in 2.5% (v/v) glutaraldehyde and 2.0% (w/v) formaldehyde in 0.1 M phosphate buffer, pH 7.4. Samples were post-fixed in 1.0% (w/v) osmium tetroxide in 0.1 M phosphate buffer at pH 7.4 for 2 hr, rinsed with deionized water three times for 5 mins each, and dehydrated in an ethanol series at room temperature. Samples were embedded in Epon resin and sectioned at a thickness of 0.5 µm and stained in 0.5% (w/v) toluidine blue O. These fixed sections were imaged using a light microscope (IX81, Olympus, Japan) and processed using ImageJ (Ferreira and Rasband, 2012). Fixed longitudinal section of young roots were prepared by fixing seedlings as described by Ursache et al., (2018). Briefly, seedlings were incubated in phosphate buffer for 1 hr and washed 3 times for 10 mins. For cell wall staining, seedlings were stained with 0.1% Calcofluor White (in ClearSee solution) for 30 mins and then washed in ClearSee for 30 mins. Root imaging was done using confocal microscopy (TCS SP8 HyD confocal microscope, Leica, Wetzlar, Germany) with an excitation window set to 405 nm and a detection window set from 425 to 475 nm.

**Stomatal staining.** Small pieces of freshly detached leaves were immersed in 1/10x (w/v) propidium iodine solution for five to ten mins and imaged on a Confocal Laser Scanning Microscopy (TCS SP8 HyD confocal microscope, Leica, Wetzlar, Germany). Propidium iodide was excited at 535 nm and the emission was collected at 570 to 630 nm. Stomata per unit area was calculated using ImageJ (Ferreira and Rasband, 2012). We counted the number of stomata on both hydroponically- and soil-grown plants and on adaxial and abaxial surfaces at the same developmental stage and treatment duration.

**Scanning electron micrographs.** For scanning electron microscopy, freshly detached leaves were mounted on the specimen stubs using double-sided tape and observed under high vacuum mode at 5.0 kV using a scanning electron microscope (JSM 6610LV, JEOL, Tokyo, Japan).
